# Interplay between cortical actin and E-cadherin dynamics regulates cell shape in the *Drosophila* embryonic epidermis

**DOI:** 10.1101/801456

**Authors:** Joshua Greig, Natalia A. Bulgakova

## Abstract

Precise regulation of cell shape is vital for building functional tissues. Here, we study the mechanisms which lead to the formation of highly elongated anisotropic epithelial cells in the *Drosophila* epidermis. We demonstrate that this cell shape is the result of two counteracting mechanisms at the cell surface: actomyosin, which inhibits cell elongation downstream of RhoA signalling, and intercellular adhesion, modulated via clathrin-mediated endocytosis of E-cadherin, which promotes cell elongation downstream of the GTPase Arf1. We show that these two mechanisms are interconnected, with RhoA signalling activity reducing Arf1 recruitment to the plasma membrane. Additionally, cell adhesion itself regulates both mechanisms: p120-catenin, a regulator of intercellular adhesion, promotes the activity of both Arf1 and RhoA. Altogether, we uncover a complex network of interactions between cell-cell adhesion, the endocytic machinery, and the actomyosin cortex, and demonstrate how this network regulates cell shape in an epithelial tissue *in vivo*.

## Introduction

The structure of all biological organisms is the product of the tissues of which they are composed. The morphogenesis of these tissues requires precise control over the shape of individual cells. In epithelial tissues, which outlines all cavities and surfaces of animal bodies, a variety of shapes and functions is observed. Cell shape is determined by mechanical properties, which define cell geometry based on intracellular and intercellular forces (Chalut and Paluch, 2016; Lecuit and Lenne, 2007). At the cell surface, mechanical properties are determined by an interplay of two factors: cortical actin and intercellular adhesion (Lecuit and Lenne, 2007; Winklbauer, 2015).

The first factor, cortical actin, is a meshwork of actin filaments crosslinked by specific cross-linking proteins and myosin motors (Chugh and Paluch, 2018). Myosin motors, such as non-muscle Myosin II (MyoII), generate contractile forces and cortical tension, which acts to minimize the contact area between cells (Blankenship et al., 2006; Clark et al., 2014). The control of cortical contractility is required to establish and maintain specific cell shapes. One of the best documented regulators of cortical contractility is the GTPase RhoA (Spiering and Hodgson, 2011). A key effector of RhoA is the enzyme Rho-Kinase (Rok), which is recruited to membranes by the activated form of RhoA, where it phosphorylates the myosin light chain, leading to activation of MyoII and an increase of actin contractility (Amano et al., 2010; Kawano et al., 1999; Kureishi et al., 1997; Leung et al., 1995).

The second factor, intercellular adhesion, is the property of one cell binding to it neighbours using specialized proteins on its surface. In epithelia, this is mediated by Adherens Junctions (AJs), with the principle component being E-cadherin (E-cad), a transmembrane protein which binds to E-cad molecules on adjacent cells (Takeichi, 1977; van Roy and Berx, 2008). Intercellular adhesion often opposes cortical tension by increasing the contact surface between cells (De Vries et al., 2004; Lecuit and Lenne, 2007), and its strength is proportional to both the levels and dynamics of E-cad at the cell surface (Foty and Steinberg, 2005; Troyanovsky et al., 2006). The latter largely relies on the processes of endocytosis and recycling, which constantly remodel AJs (Kowalczyk and Nanes, 2012).

The p120-catenin (p120ctn) protein family are the key regulators of E-cad endocytosis in mammalian cells, through directly binding the juxtamembrane domain of E-cad (Daniel and Reynolds, 1995; Garrett et al., 2017; Ireton et al., 2002; Oas et al., 2013; Shibamoto et al., 1995; Yap et al., 1998; Yu et al., 2016). This catenin family is represented by a single gene in invertebrates, such as *Drosophila*, whereas p120ctn has 7 members in humans with different expression patterns and functional requirements, reflecting an increase in tissue types and complexity (Carnahan et al., 2010; Gul et al., 2017; Hatzfeld, 2005). Most studies have focused on the founding family member, p120ctn, however other members, δ-catenin and ARVCF, seem to have similar functions (Davis et al., 2003). In mammalian cells, p120ctn is required to maintain E-cad at the plasma membrane: uncoupling p120ctn from E-cad or reducing expression results in complete internalization of E-cad (Davis et al., 2003; Ireton et al., 2002; Ishiyama et al., 2010), as the binding of p120ctn to E-cad conceals endocytosis triggering motifs (Nanes et al., 2012; Reynolds, 2007). This model of p120ctn activity has recently been augmented in mammalian cells, when it was found that p120ctn can also promote endocytosis of E-cad through interaction with Numb (Sato et al., 2011).

By contrast, in *Drosophila* and *C. elegans*, p120ctn was thought to play only a supporting role in adhesion as genetic ablation failed to replicate the effects observed in mammalian systems (Myster et al., 2003; Pacquelet et al., 2003; Pettitt et al., 2003). This was thought to be due to the greater similarity of invertebrate p120ctn to mammalian δ-catenin, ablation of which is similarly viable in mice (Carnahan et al., 2010; Israely et al., 2004). However, δ-catenin expression is restricted to neural and neuroendocrine tissues (Ho et al., 2000), which is likely to explain the mildness of knockout phenotypes, whereas invertebrate p120ctn is broadly expressed in both epithelia and neurons (Myster et al., 2003), suggesting potential functional similarity with mammalian p120ctn which shares the broad expression pattern (Davis et al., 2003). Furthermore, it has recently been reported that *Drosophila* p120ctn is required to stabilize E-cad in the pupal wing (Iyer et al., 2019) and promotes the endocytosis and recycling of E-cad in the embryo and larval wing discs (Bulgakova and Brown, 2016). This indicates an evolutionary conservation of p120ctn function, where depending upon the context, p120ctn either inhibits or promotes E-cad endocytosis.

Another important protein family regulating E-cad endocytosis is the Arf GTPases, which recruit coat proteins to facilitate the intracellular trafficking of vesicles (Donaldson and Jackson, 2011). The first member of the family, Arf1, is classically viewed as Golgi resident and responsible for anterograde transport from the Golgi to the plasma membrane (Donaldson and Jackson, 2011; McMahon and Boucrot, 2011). Recently, however, Arf1 was detected at the plasma membrane, where it participates in trafficking by co-operating with Arf6-dependent endocytosis (Humphreys et al., 2013; Padovani et al., 2014). In *Drosophila*, Arf1 is required for facilitating endocytosis in the early syncytial embryo (Humphreys et al., 2012; Lee and Harris, 2013; Rodrigues et al., 2016). Indeed, Arf1 interacts with E-cad and another component of AJs, Par-3 (Shao et al., 2010; Toret et al., 2014).

Cortical actin and intercellular adhesion do not exist in isolation but are intimately interconnected. Intracellularly E-cad interacts with other catenin proteins: at its distal C-terminus E-cad binds β-catenin, with α-catenin binding β-catenin and actin, thus directly linking E-cad to cortical actin (Ozawa et al., 1990; Shapiro and Weis, 2009). In addition to direct linkage, cortical actin and intercellular adhesion share common regulators. In in mammalian epithelial cells, RhoA localizes to AJs, where E-cad adhesion complexes create local zones of active RhoA by recruiting Ect2 GEF through α-catenin (Priya et al., 2013; Ratheesh et al., 2012). In these cells, p120ctn has a context-dependent role in RhoA regulation: it can directly inhibit RhoA, indirectly via p190RhoGAP, direct its spatiotemporal activity, or activate it (Anastasiadis et al., 2000; Derksen and van de Ven, 2017; Lang et al., 2014; Taulet et al., 2009; Yu et al., 2016; Zebda et al., 2013). The role of p120ctn in the RhoA pathway in *Drosophila* is unclear (Fox et al., 2005; Magie et al., 2002). However, RhoA itself regulates E-cad and is required for establishment of E-cad-mediated adhesion (Braga et al., 1997). In *Drosophila*, RhoA promotes the regulated endocytosis of E-cad by Dia and AP2 (Levayer et al., 2011b). Conversely, RhoA activity antagonises E-cad endocytosis in the early embryo (Lee and Harris, 2013), indicating a context-dependent role of RhoA in E-cad endocytosis. Finally, E-cad endocytosis can direct actin remodelling (Hunter et al., 2015). However, how this interplay between cortical actin and intercellular adhesion produces cell shape has yet to be fully elucidated.

Here, we investigated the interplay between cortical actin and E-cad-mediated adhesion in the highly elongated cells of the late embryonic epidermis in *Drosophila.* We demonstrate that this cellular elongation is accompanied by a reciprocal anisotropy of cortical tension relative to E-cad adhesion. We determined that the prototypical member of the p120ctn family is directly involved in shaping these cells. p120ctn achieves this by both influencing the endocytosis of E-cad and by modulating cortical tension. We characterised the mechanism of this p120ctn dual function and found that it is dependent on interactions with two GTPase signalling pathways: RhoA to increase cortical tension and inhibit endocytosis, and Arf1 to promote endocytosis and enable the remodelling of the AJs. Finally, we show that the interplay between these two GTPase pathways downstream of p120ctn directly participates in shaping the morphology of cells *in vivo*.

## Results

### 1) Anisotropy of cortical actin and adhesion in the epidermis of stage 15 Drosophila embryos

The epidermal cells of stage 15 *Drosophila* embryos have a distinct rectangular morphology in their apical plane (Fig. 1A-C). These cells are elongated in the direction of the dorsal-ventral axis of the embryo and have the average aspect ratio of 6.1 ± 1.3 (ratio between lengths of the long and short cell axes, mean ± SD, Fig. 1D). Consistent with the elongated shape, these cells have two distinct types of cell border: long borders, which are orthogonal to the anterior-posterior axis of the embryo (AP borders), and short borders, which are orthogonal to the dorsal-ventral axis (DV borders, Fig. 1B-C).

**Figure 1.**
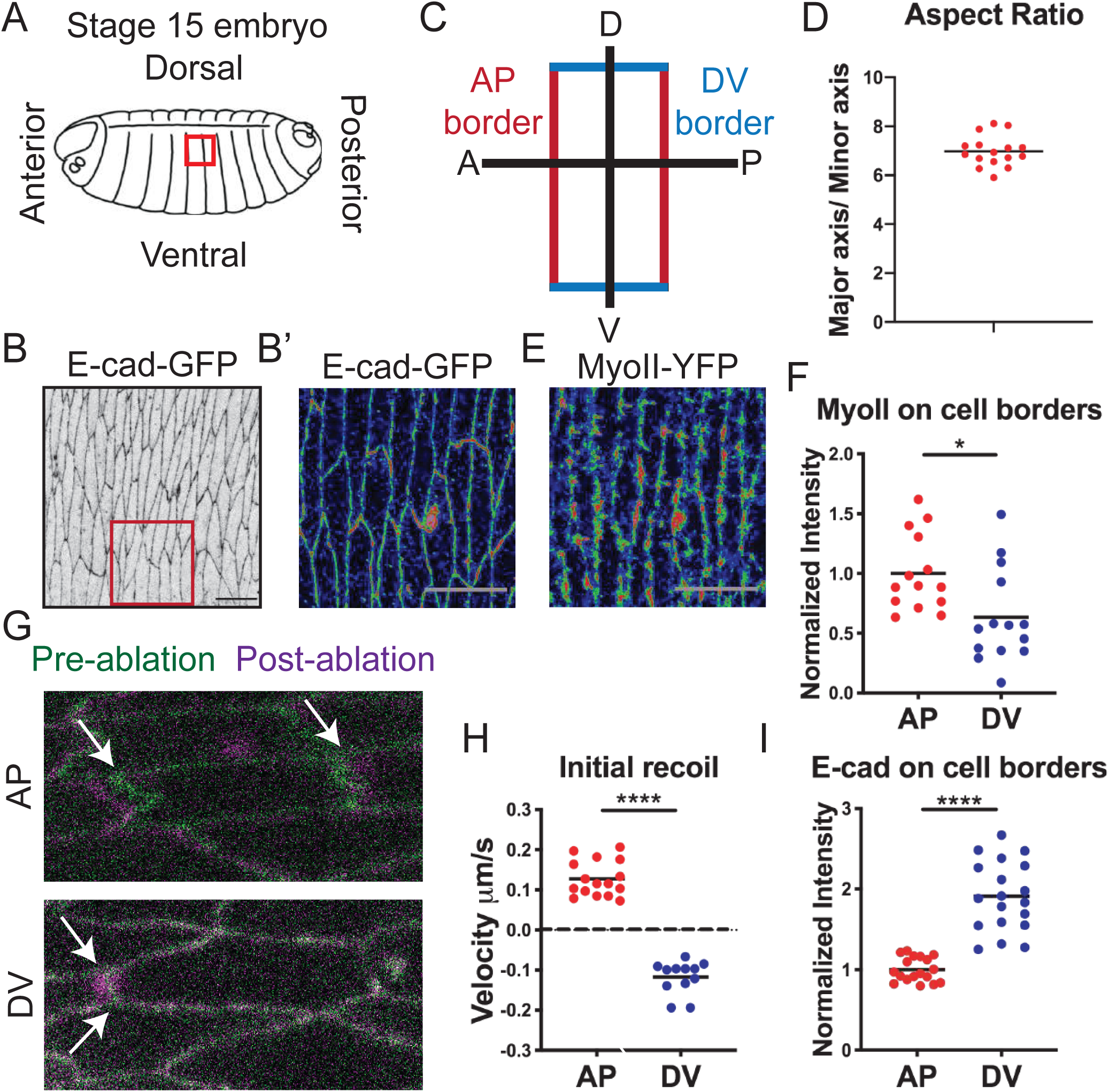
Cells of the stage 15 *Drosophila* embryonic epidermis have anisotropic cell surface adhesion, actomyosin, and cortical tension. **A-D.** Overview of the stage 15 *Drosophila* embryo. (A) A cartoon sketch of a stage 15 *Drosophila* embryo, area indicted by red box is the dorsolateral epidermis region imaged in this study. (**B**) Apical view of the epidermis highlighted in the red box (A). Cells are outlined by E-cad-GFP. (**B’**) Magnified image indicated in the box of (B) displayed using rainbow pixel intensity spectrum to highlight asymmetry of distribution, scale bar is 10 µm. (**C**) A schematic of the two cell borders present in these cells: the longer Anterior-Posterior (AP, red) and the short Dorsal-Ventral (DV, blue). (**D**) Aspect ratio, which is the length of the cell in the long AP axis divided by the length in the short DV, of the cells in embryos expressing only E-cad-GFP in otherwise wild-type background. (**E**) Non-muscle Myosin II (MyoII) tagged with YFP in the same plane as E-cad (B) and displayed using rainbow pixel intensity spectrum. (**F-G**) Amounts of MyoII-YFP (F) and E-cad-GFP (G) at the AP and DV cell borders in embryos expressing only either MyoII-YFP (F) or E-cad-GFP (G) in otherwise wild-type background. (**H**) Examples of laser ablation of an AP border (upper panel) and a DV border (lower panel) in dorsolateral epidermal cells of embryos expressing E-cad-GFP to visualize cell borders, in otherwise wild type background. Image is an overlay of pre-ablation (green) and post-ablation (magenta). The arrows indicate the connected vertices of the ablated membrane used to measure initial recoil. Note, the recoil clearly seen in ablation of the AP border is absent in ablation of DV. Statistical difference between cell borders measured by two-way ANOVA. *, P < 0.05; ***, P < 0.001. 10-20 embryos per genotype were quantified.

These AP and DV cell borders differ in their actin cortex, with MyoII enriched at the AP borders in these cells (AP:DV≅2:1, Fig 1E-F), consistent with previous reports (Bulgakova et al., 2013; Simoes et al., 2010). As the accumulation of MyoII has been linked to cortical tension (Priya et al., 2015; Scarpa et al., 2018; Yu and Fernandez-Gonzalez, 2016), we compared tension between the AP and DV borders using microablation and measured the initial recoil as it is proportional to the tension in the system rather than other variables (Liang et al., 2016; Mao et al., 2013). We used an E-cad tagged at its endogenous locus (E-cad-GFP, Huang et al., 2009) to label cell outlines and quantify the initial recoil (Fig. 1G-H). We found that while the initial recoil was positive for the AP borders, showing that they are under tension, it was negative for the DV borders, suggesting that they are under compression (Fig. 1H).

Further to the difference in MyoII distribution and tension properties, the AP and DV borders display a substantial difference in the distribution of intercellular adhesion components, specifically, an asymmetry in the levels and dynamics of E-cad. In these differentiated cells E-cad localizes in a narrow continuous band of fully formed mature AJs (Adams et al., 1996; Tepass and Hartenstein, 1994). E-cad-GFP localized asymmetrically with a 1:2 (AP:DV) ratio between the two cell borders (Fig. 1I, Table S1, and Bulgakova et al., 2013). This asymmetry is due to an accumulation at the DV borders of a specific pool of E-cad, which is highly dynamic due to its endocytic trafficking (Bulgakova et al., 2013). Overall, the elongated shape of these epidermal cells coincides with anisotropies in both cortical actin and tension, and the levels and dynamics of adhesion complexes. Therefore, we investigated how the interplay between cortical tension and adhesion at the cell surface produces cell shape, and how the interaction between these two processes is mediated.

### 2) p120ctn influences cell shape, RhoA signalling and cortical tension

Members of the p120ctn family regulate both the actin cytoskeleton and cadherin trafficking and are thus good candidates to mediate the interplay between cortical tension and cell adhesion. In this vein, we investigated if the function of p120ctn in *Drosophila*, which have only one member of the family, affected the shape of cells. We created a p120ctn overexpression construct under the control of a *UAS* promoter (*UAS*::p120ctn) and expressed it in the posterior half of each embryonic segment using the *engrailed*::GAL4 (*en*::GAL4) driver while marking the cells using *UAS::*CD8-Cherry (Fig. 2A). To exclude potential differences between the compartments we used an external control: an *engrailed* compartment expressing two copies of *UAS::*CD8-Cherry to balance the Gal4:UAS ratio (Fig. 2A). We complemented this analysis by using a *p120ctn* mutant to examine the effect of p120ctn loss (Fig. 2A, and Bulgakova and Brown, 2016). The cells expressing *UAS*::p120ctn appeared distorted and had a smaller aspect ratio (p<0.0001, Fig. 2A-B), indicative of cell rounding. By contrast the loss of p120ctn did not change the aspect ratio (p=0.27, Fig. 2A,C), consistent with the previous report (Bulgakova and Brown, 2016). Therefore, increased p120ctn levels have a distinct effect on the shapes of cells in the *Drosophila* epidermis, altering their apparent elongation and anisotropy.

**Figure 2.**
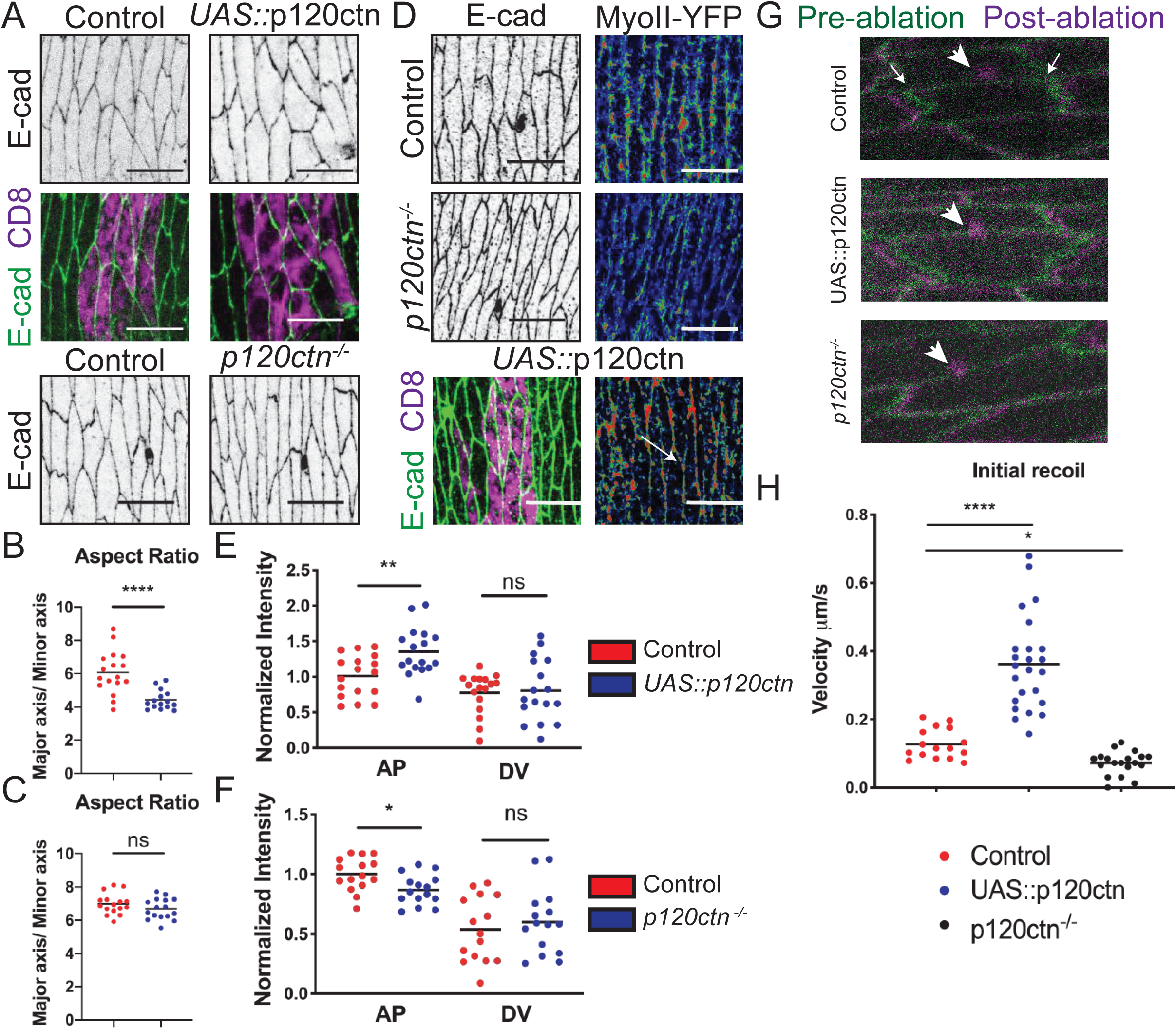
p120ctn levels affect cell shape and cortical actin, resulting in concomitant changes in cortical tension. (**A)** Top and middle: Apical views of epidermis expressing *UAS::*CD8-Cherry (left) and co-expressing it with *UAS::*p120ctn (right) using *en:*:GAL4 with E-cad-GFP (green, top; grey middle) and *UAS::*CD8-Cherry (magenta, top). *UAS::*CD8-Cherry was used to mark cells expressing transgenes. Bottom: Control and *p120ctn^-/-^* mutant epidermal cells marked by E-cad-GFP (grey). (**B**) Aspect ratio of the control (*UAS::*CD8-Cherry) and *UAS::*p120ctn expressing cells. (**C**) Aspect ratio of the wide-type control and *p120ctn^-/-^* mutant cells. (**D**) The epidermis of control, *p120ctn^-/-^* mutant and p120ctn overexpression (*UAS::*p120ctn) embryos visualized with anti-E-cad antibody (grey, left and middle; green, bottom), *UAS*::CD8-mCherry (magenta, bottom image, marks cells expressing *UAS::*p120ctn), and Myosin II (MyoII) tagged with YFP (rainbow pixel intensity spectrum, right, membrane indicated by arrow). (**E-F**) Amounts of MyoII-YFP in p120ctn overexpressing embryos (E) and in the *p120ctn^-/-^*mutant (F). All membrane intensity data are in Table S1. Scale bars – 10µm. (**G**) Laser ablation of AP cell borders in control (top), p120ctn overexpression (middle), and *p120ctn^-/-^* mutant (bottom). Images are an overlay of pre-ablation (green) and post-ablation (magenta). The small arrows indicate the connected vertices of the ablated membrane used to measure initial recoil. The large arrows represent the position of microablation, for which a small spot of discolouration can be observed. (**H**) The initial recoil velocity of the membranes in the three conditions. Statistical difference between cell borders measured by two-way ANOVA. For laser ablation statistical analysis is a two-tailed students t-test with Welch’s correction. All best-fit and membrane intensity data are in Table S1. *, P < 0.05; **, P < 0.01 ****, P < 0.0001. 10-20 embryos per genotype.

A well-established regulator of cell shape is cortical actin, which is highly anisotropic in epidermal cells and is regulated by RhoA-signalling. As p120ctn family members have been reported to both positively and negatively regulate RhoA-signalling, we examined the interaction between p120ctn and RhoA in our system. We did so by examining its downstream effectors: the enzyme Rok and its target MyoII. p120ctn overexpression increased the amount of MyoII-YFP at the AP borders (p=0.006, Fig. 2D-E). Conversely, in the *p120ctn* mutant embryos MyoII-YFP was reduced specifically at the AP borders (p=0.01, Fig. 2D,F). The amount of MyoII-YFP at the DV borders were not affected in both cases (p=0.47 and p=0.45, Fig. 2E-F). We complemented these experiments by using a tagged kinase-dead variant of Rok (Venus-Rok^K116A^, referred to as Rok-Venus, Simoes et al., 2010), previously used as a readout of Rok localization and activity (Bulgakova et al., 2013; Simoes et al., 2010). Rok-Venus had the same localization and was affected in the same way as MyoII-YFP (compare Fig. S1 with Fig. 2E-F). p120ctn loss abolished Rok-Venus asymmetry due to a reduction at the AP borders, and overexpression increased Rok-Venus at the AP borders (p=0.013 and p=0.0043, respectively, Fig. S1). These results indicate that p120ctn activates RhoA signalling specifically at the AP borders in a dose-dependent manner in these epidermal cells.

Due to this positive correlation between p120ctn levels and RhoA signalling on the long AP cell border, we decided to directly measure the cortical tension at the AP borders in *p120ctn* mutant and overexpressing cells (Fig. 2G-H). The overexpression of p120ctn increased the mean initial recoil (0.36 µm/sec in comparison to 0.13 µm/sec, p<0.0001, Fig. 2G-H). Conversely, in *p120ctn* mutant cells the recoil was decreased to a mean value of 0.07 µm/sec (p=0.022, Fig. 2G-H). This demonstrates that cortical tension correlates with p120ctn levels. Overall, the changes in p120ctn levels altered the shape of the epidermal cells, with the levels of p120ctn positively correlating with the activity of RhoA signalling and cortical tension at the AP borders. The increased tension at the AP borders following p120ctn overexpression is also consistent with the rounded cell phenotype. The DV borders displayed no change in the activity of RhoA signalling, indicating an anisotropic positive action of p120ctn on RhoA signalling.

### 3) p120ctn regulates E-cad amounts and dynamics within adhesion sites

The other factor which contributes to cell shape is intercellular adhesion which is mediated by E-cad in epithelial cells. In our system this intercellular adhesion is also highly anisotropic with an opposite pattern to the cortical actin (Bulgakova et al., 2013). p120ctn binds to the intracellular domain of E-cad, enabling it to regulate the endocytosis of E-cad (Bulgakova and Brown, 2016; Iyer et al., 2019; Nanes et al., 2012; Reynolds, 2007; Sato et al., 2011). First, we determined that p120ctn in *Drosophila* co-localizes with E-cad. To test this, we used ubiquitously expressed p120ctn tagged with GFP (*Ubi*::p120ctn-GFP) and found that it localized to both AP and DV borders with an enrichment at the DV, mimicking the localization of E-cad-GFP (r = 0.868, p <0.00001, Fig. 3A-B). Note that in this system, the antibody against the N-terminus of p120ctn does not work and fails to reproduce the localization of the full-length GFP tagged p120ctn (Fig. S2). Given this colocalization, we then examined if changes in E-cad levels were observed when altering the levels of p120ctn. In cells overexpressing p120ctn we detected a reduced asymmetry of E-cad-GFP due to an increase in levels specifically at the AP but not DV borders (p=0.0002 and p=0.27, Fig. 3C-D, and see Fig. 2A). In contrast, the loss of p120ctn resulted in an isotropic decrease in E-cad-GFP amounts at both AP and DV borders by approximately 15% (p=0.008 and p=0.035, Fig. 3E-F, see also Fig. 2A).

**Figure 3.**
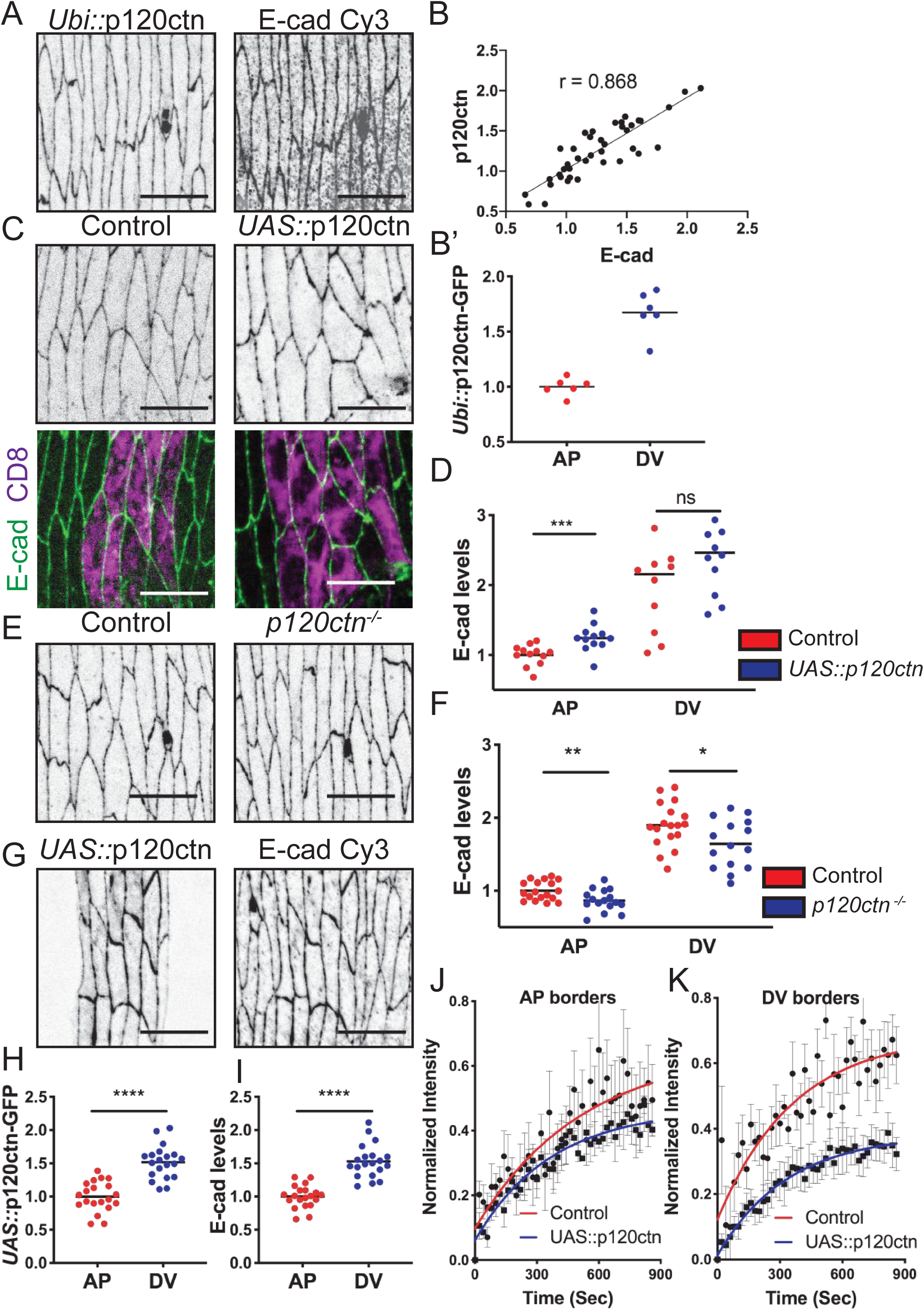
p120ctn regulates both the levels and dynamics of E-cad. **(A)** Apical view of epidermal cells expressing a *Ubi::*p120ctn-GFP transgene, cell borders visualised by transgene itself (grey, left) and stained with E-cad antibody (grey, right). (**B-B’**) Quantification of the localization of the *Ubi::*p120ctn-GFP transgene in the epidermal cells. (B) Pearson’s correlation of the signal intensities between E-cad and *Ubi::*p120ctn-GFP. (B’) The localization of the *Ubi::*p120ctn-GFP transgene on the two cell borders. (**C-D**) Effect on E-cad of overexpressing p120ctn using *UAS*::p120ctn construct. Apical views (C) and the membrane E-cad levels (D) in cells expressing E-cad-GFP (C, grey, top; green bottom) and *UAS*::CD8-mCherry (C, control, magenta, left) or *UAS*::p120ctn co-expressing a single copy of *UAS*::CD8-mCherry (C, magenta, right). (**E-F**) Apical views (E) and the membrane E-cad levels (F) in epidermis of control and *p120ctn^-/-^*mutant cells both expressing endogenous E-cad-GFP. (**G-I**) Localization of E-cad and p120ctn in embryos expressing a *UAS*::p120ctn-GFP. (G) *UAS*::p120ctn-GFP expressed in engrailed compartment cells using the *engrailed*::GAL4 driver. (G) The same cells stained for E-cad to compare with internal control cells adjacent to engrailed stripe. (H-I) The localization of *UAS*::p120ctn-GFP (H) and E-cad (I) in the *UAS*::p120ctn-GFP expressing cells between the two borders. (**J-K**) Dynamics of E-cad-GFP measured by FRAP in control and *UAS::*p120ctn cells at the AP (J) and DV (K) borders. Average recovery curves (mean ± s.e.m.) and the best-fit curves (solid lines) are shown. All best-fit and membrane intensity data are in Table S1. Statistical difference between cell borders was measured by two-way ANOVA. For comparison of the difference between cell borders a two-tailed students t-test with Welch’s correction. Scale bars – 10 µm. *, p < 0.05, **, p < 0.01 ***, p < 0.001; ****, p < 0.0001. 10-20 embryos per genotype were used for quantifications; 8-10 embryos – for FRAP.

Next, we examined how this increase in the E-cad amount at the AP borders correlated with p120ctn localization. We overexpressed a tagged p120ctn-GFP (*UAS::*p120ctn-GFP) driven by *en::*GAL4 to compare p120ctn and E-cad localization in the same cell. We detected an AP:DV ratio of 2:3 for both E-cad and p120ctn (Fig. 3G-I, Fig. S2). The lower than usual AP:DV ratio of E-cad was similar to that observed when untagged p120ctn was overexpressed. As this E-cad distribution was identical to the distribution of p120ctn itself, we concluded that additional E-cad molecules are recruited as a complex with p120ctn.

The strength of cell-cell adhesion as well as the amount of adhesion complexes at the cell surface are regulated by endocytosis, and p120ctn has been shown to inhibit and promote E-cad endocytosis in mammalian and *Drosophila* cells, likely reflecting context-dependent functions (Bulgakova and Brown, 2016; Ireton et al., 2002; Nanes et al., 2012; Sato et al., 2011; Xiao et al., 2003). Therefore, we decided to explore whether the changes in the levels of E-cad are accompanied by changes in its dynamics. We used Fluorescence Recovery After Photobleaching (FRAP), which reveals the stable fraction of the protein which does not exchange within the sites of cell-cell adhesion on the time-scale of experiment, and the mobile fraction. E-cad-GFP mobile fraction is 70% for DV and 50% for the AP borders in control, with 30% and 50% of protein being stable, respectively (Fig. 3J-K). The recovery curves which describe the dynamics of E-cad-GFP mobile fraction were best-fit by a bi-exponential model (Table S1, and Bulgakova et al., 2013), with the fast and slow components attributed to diffusion and endocytic recycling, respectively (Bulgakova et al., 2013; Iyer et al., 2019; Lippincott-Schwartz et al., 2003).

Measuring the dynamics of E-cad-GFP in p120ctn overexpressing cells we discovered that it was less dynamic at both border types (Fig. 3J-K). The mobile fractions were approximately 40% and 30% for AP and DV borders respectively, which resulted in an increase of the immobile fraction at both borders to 60% and 70% relative to control (p=0.0023 and p<0.0001 Table S1). This is similar to changes in the dynamics of E-cad-GFP in *p120ctn* mutants, where an increase in the immobile E-cad-GFP fraction is also observed (Fig. S2, and Bulgakova and Brown, 2016). Therefore, while p120ctn overexpression elevates E-cad level at the AP borders, and its loss uniformly reduces it, both lead to stabilization of E-cad-GFP at cell junctions. Next, we sought to determine the mechanism of this stabilization and how these changes in E-cad dynamics contribute to cell shape changes caused by elevated p120ctn levels.

### 4) p120ctn and RhoA regulate E-cad via clathrin-mediated endocytosis

To ascertain if the stabilization of E-cad-GFP by altered p120ctn levels was due to an impairment of endocytosis, we examined clathrin, which is the main coat protein involved in clathrin-mediated endocytosis. We used the Clathrin Light Chain (CLC) tagged with GFP (*UAS*::CLC-GFP, Loerke et al., 2005; Wu et al., 2001), to monitor clathrin behaviour in the plane of AJs by performing FRAP. CLC tagged with GFP incorporates functionally into clathrin-coated pits without interfering with basic clathrin functions (Chang et al., 2002; Gaidarov et al., 1999; Kochubey et al., 2006). CLC-GFP expressed using the *en*::GAL4 driver was found in spots on the plasma membrane in the plane of AJs and in the cytoplasm (Fig. 4A), a localization consistent with its function (Kaksonen and Roux, 2018). p120ctn overexpression reduced the recovery of the CLC-GFP, with a reduction in the mobile fraction by 30% relatively to control (p<0.0001, Fig. 4B). A similar reduction in CLC-GFP mobile fraction by 25% was observed in *p120ctn* mutants (p<0.0001, Fig. 4C). These reflect the changes in the mobile fraction of E-cad (Fig. 3J-K), suggesting that CLC-GFP recovery is a valid proxy for E-cad dynamics in these cells.

**Figure 4.**
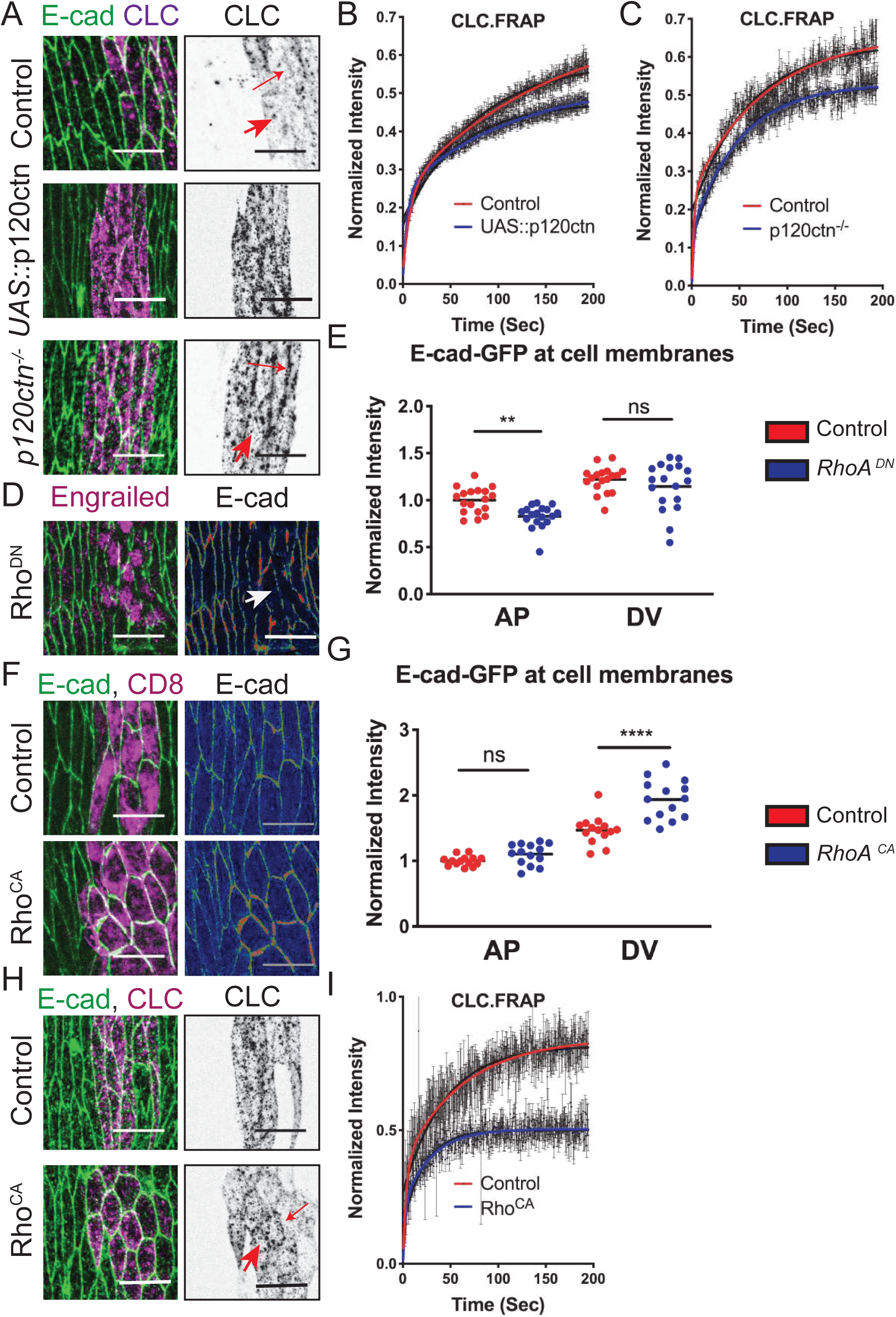
p120ctn and RhoA change the dynamics of the clathrin mediated endocytosis. (**A-C**). The localization and dynamics of the clathrin light chain (*UAS*::CLC-GFP) in control, embryos overexpressing p120ctn (*UAS*::p120ctn), and *p120ctn^-/-^* mutant embryos. (A) Distinct puncta (spots, magenta in left and black in right images, arrows) are observed at the membrane and in the cytoplasm. Cell outlines are visualized by anti-E-cad antibody (green, left). FRAP of *UAS*::CLC-GFP using a rectangular region in the plain of AJs in embryos overexpressing p120ctn (*UAS*::p120ctn) (B) and *p120ctn^-/-^*mutant (C). Average recovery curves (mean ± s.e.m.) and the best-fit curves (solid lines) are shown. (**D-G**). Localization of E-cad in the epidermis following downregulation (the induction of Rho^DN^ expression for 4 hours, membrane indicated by arrow, D-E) or upregulation (expression of Rho^CA^, F-G) of RhoA signalling. Top images in **F** are the corresponding control expressing *UAS*::CD8-Cherry alone. Cell borders are marked by E-cad-GFP (D and F, rainbow spectrum, right; green, left). Cells expressing transgenes are marked with an antibody for Engrailed (D) or by *UAS::*CD8-Cherry (F). Levels of the E-cad-GFP at the membrane of the Rho^DN^ (E) or Rho^CA^ (G) expressing cells in comparison to internal control (E) or external controls (G). (**H-I**). Localization of clathrin (*UAS*::CLC-GFP, grey, right; magenta, left) in cells co-expressing Rho^CA^ (bottom) and corresponding control co-expressing *UAS*::CD8-Cherry (top). Cell borders are visualized by anti-E-cad antibody (green, left, arrow indicates membrane localized puncta). (**I).** FRAP of *UAS*::CLC-GFP in Rho^CA^ expressing cells. Average recovery curves (mean ± s.e.m.) and the best-fit curves (solid lines) are shown. All best-fit and membrane intensity data are in Table S1. Scale bars – 10 µm. Statistical analysis is a two-tailed students t-test with Welch’s correction. *, p < 0.05; **, p < 0.01 ****, p < 0.0001. 10-20 embryos per genotype were used for quantifications; 8-10 embryos – for FRAP.

As p120ctn overexpression resulted in both an increase of E-cad levels and anisotropic activation of RhoA signalling, we asked whether this change in the RhoA signalling was responsible for the changes in E-cad, for instance by modulating its endocytosis. We sought to mimic the RhoA phenotype in *p120ctn* mutants by directly inhibiting the RhoA pathway using a dominant negative (Rho^DN^) construct. Rho^DN^, expressed constitutively using *en::GAL4*, resulted in a complete loss of E-cad at the membrane by stage 15 of embryogenesis, therefore, we acutely induced the expression of the Rho^DN^ using ubiquitously expressed temperature sensitive GAL80^ts^ and a temperature shift for 4 hours at 12 hours after egg laying (Pilauri et al., 2005). This acute downregulation of RhoA signalling reduced the amounts of E-cad-GFP at the AP but not DV borders (p=0.0078 and p=0.37, Fig. 4D-E). Note that in this case E-cad asymmetry was reduced in control cells, likely due to the effects of temperature shift during induction of expression. In a complementary experiment, we expressed a constitutively-active Rho^CA^ to elevate RhoA signalling directly (Fig. 4F). The expression of Rho^CA^ increased the amounts of E-cad-GFP at the cell surface, specifically at the DV but not the AP borders (p<0.0001 and p=0.34, Fig. 4G), which is consistent with an ectopic activation of RhoA signalling at the DV borders in this case. We then asked if this increase in E-cad could be explained by the stabilization of E-cad-GFP at the plasma membrane by elevated RhoA signalling. To this end, we measured the dynamics of clathrin in the plain of AJs using FRAP as a proxy for E-cad endocytosis. Indeed, the mobile fraction of CLC-GFP was reduced by 40% in comparison to control (p<0.0001, Fig. 4H-I). In cells overexpressing p120ctn, increase of E-cad at the AP borders was accompanied by increased activation of RhoA signalling. Overall, these data suggest that p120ctn leads to activation of RhoA signalling at the AP borders, which elevates E-cad most likely by preventing its endocytosis at these borders.

### 5) The localization of the GTPase Arf1 at the cell plasma membrane depends on p120ctn and RhoA, and promotes clathrin-mediated endocytosis

Elevated RhoA signalling resulted in E-cad stabilization at the cell surface and inhibited clathrin-mediated endocytosis, however E-cad was also stabilized when RhoA signalling was downregulated in *p120ctn* mutants. Therefore, we next sought to identify molecules which were responsible for this E-cad stabilization in *p120ctn* mutants. The GTPase Arf1 has been reported to interact with E-cad and other AJ components (Shao et al., 2010; Toret et al., 2014). Therefore, we examined if Arf1 acts downstream of p120ctn using a transgenic GFP-tagged variant of Arf1 (*UAS*::Arf1-GFP, Lee and Harris, 2013) driven by *en::*GAL4. *UAS*::Arf1-GFP has reduced affinity for ArfGAPs and ArfGEFs, and nucleotide exchange rate (Jian et al., 2010), which allowed us to study Arf1 without hyperactivating the pathway.

*UAS*::Arf1-GFP localized to both the Golgi apparatus and plasma membrane (Fig. 5A, Fig. S3), consistent with previous reports (Lee and Harris, 2013; Shao et al., 2010). The Golgi-resident Arf1 appeared in large and distinct Arf1-GFP puncta throughout the cytoplasm including in close proximity to the plasma membrane (Fig. 5A, Fig. S3). As we were interested in the localization of Arf1 at the plasma membrane, we excluded the puncta from our analysis (see Materials and Methods). In the control, Arf1-GFP localization at the plasma membrane was uniform (p=0.36, Fig. 5C). The loss of p120ctn resulted in a uniform decrease in the amount of Arf1-GFP at both the AP and DV borders (p<0.0001 and p<0.0001, Fig. 5B-C), suggesting that p120ctn promotes Arf1 localization.

**Figure 5.**
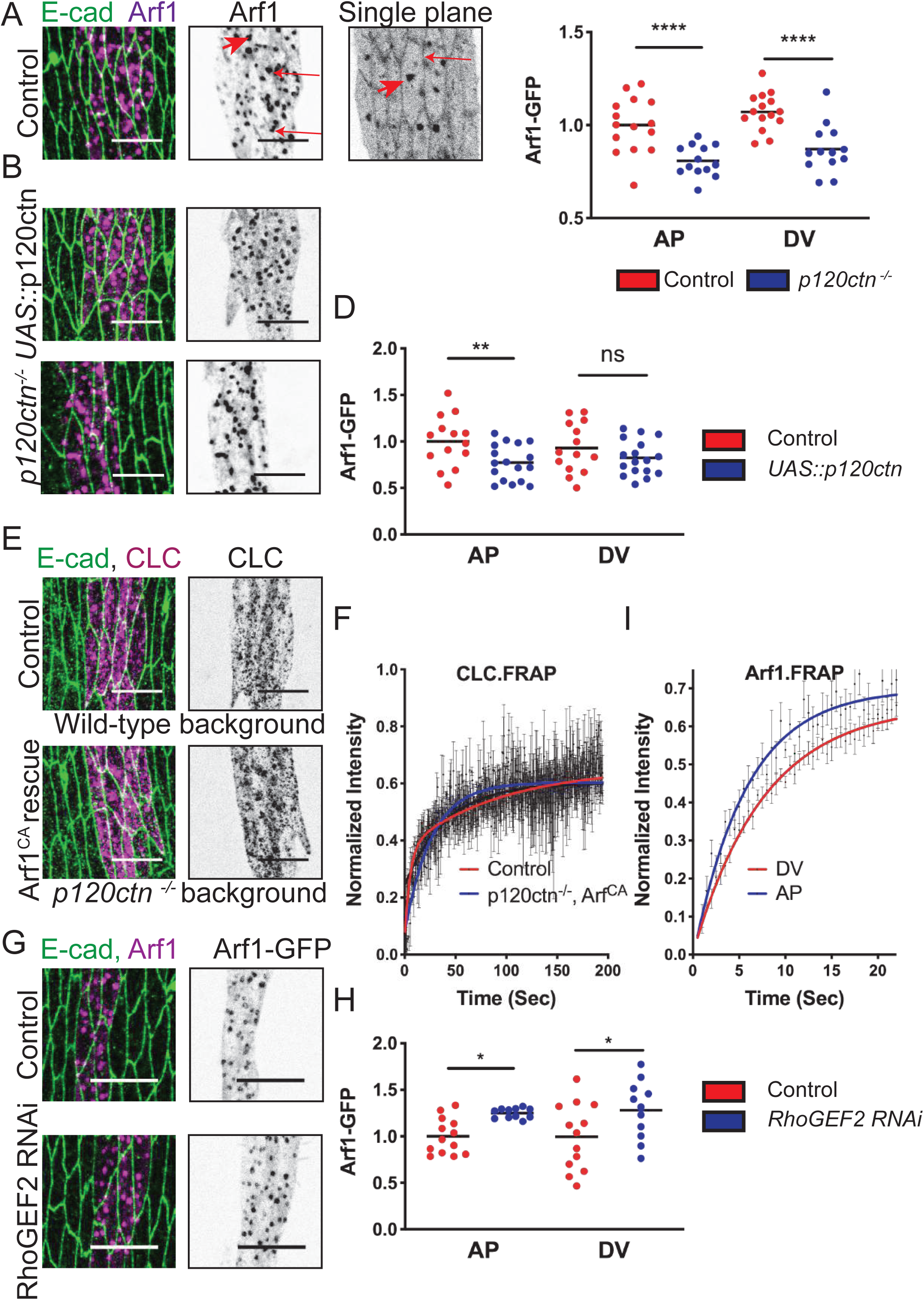
Arf1 signalling is downstream of p120ctn and promotes clathrin mediated endocytosis, but is inhibited by RhoA signalling. **(A)** Apical views of the epidermis expressing *UAS*::Arf1-GFP (Arf1, magenta, left; grey, middle and right) with *en*::Gal4. Cell outlines are visualized with anti-E-cad antibody (green, left). The large UAS::Arf1-GFP puncta in the cytoplasm (large arrow in A) mark the Golgi. The presence of the secondary Arf1-GFP population at the cell surface is revealed with a single plane section in the centre of the AJ (grey, right, small arrow), and is less prominent in an average projection spanning the whole depth of the AJ (middle). (**B**) Apical views of the epidermis expressing *UAS*::Arf1-GFP in cells overexpressing p120ctn (*UAS*::p120ctn) and *p120ctn^-/-^*mutant. (**C-D**) The amount of *UAS*::Arf1-GFP at the plasma membrane between a control and *p120ctn^-/-^* mutant (C), and between a control and p120ctn overexpressing embryos (D). **(E)** Localization of *UAS*::CLC-GFP (grey, right; magenta, left) in control, expressing *UAS*::CLC-GFP alone (top), and in embryos expressing a constitutively active Arf1 (Arf1^CA^) in a *p120ctn^-/-^* mutant genetic background (bottom). Cell borders were visualized with anti-E-cad antibody (green, left). (**F**) FRAP of *UAS*::CLC-GFP in the plane of AJs in the Arf1^CA^; *p120ctn^-/-^*embryos. Average recovery curves (mean ± s.e.m.) and the best-fit curves (solid lines) are shown. (**G-H).** The apical view (G) and amount (H) of *UAS::*Arf1-GFP (grey, right; magenta, left) in control and cells expressing RhoGEF2 RNAi. (**I**). Dynamics of Arf1-GFP in control cells using FRAP. Statistical analysis between cell borders measured by two-way ANOVA. *, p < 0.05; ***, p < 0.001. All best-fit and membrane intensity data are in Table S1. 10-20 embryos per genotype were used for quantifications; 8-10 embryos – for FRAP.

Considering this role of p120ctn in Arf1 localization and the known function of Arf1 in intracellular trafficking, we decided to test if the reduction in Arf1 activity is responsible for the CLC-GFP stabilization in *p120ctn* mutants. Therefore, we expressed a constitutively active Arf1 construct (Arf1^CA^) in *p120ctn* mutant embryos and measured the FRAP of CLC-GFP. Measuring the dynamics of clathrin revealed that the mobile fraction which recovered was no longer different from the wild-type control (p=0.19, Fig. 5E-F). We concluded that the expression of Arf1^CA^ rescues the clathrin dynamics in the *p120ctn* mutant. This is consistent with Arf1 acting downstream of p120ctn, providing a link between the p120ctn–E-cad complex and the clathrin-mediated endocytic machinery.

Curiously, the overexpression of p120ctn also resulted in a reduction of Arf1-GFP at the AP borders (p=0.02, Fig. 5B,D). Although the reduction of Arf1 at the DV borders was not significant (p=0.37), its distribution remained uniform (p=0.18). Due to evidence that GTPases often regulate each other (Baschieri and Farhan, 2012; Singh et al., 2017), we asked whether this reduction was a consequence of p120ctn elevating RhoA signalling. We measured the membrane levels of Arf1-GFP in cells expressing RhoGEF2-RNAi or *UAS*::CD8-Cherry (Fig. 5G-H). Expression of this RNAi against RhoGEF2 reduces Rok amounts by 20% specifically at the AP borders (Bulgakova et al., 2013). The downregulation of RhoGEF2 resulted in an increase in the amount of Arf1-GFP at both AP and DV borders (p=0.049 and p=0.022, respectively, Fig. 5H), demonstrating that RhoA signalling negatively regulated Arf1 localization to the plasma membrane. In a complementary experiment, we measured the membrane localization of MyoII-YFP upon the upregulation of Arf1 signalling using a constitutively active variant of Arf1 (Arf1^CA^, Fig. S3). MyoII-YFP localization was indistinguishable between cells expressing Arf1^CA^ and control cells (Fig. S3), suggesting that RhoA signalling in the embryonic epidermis is independent of Arf1 function. Therefore, we concluded that the reason Arf1 was reduced at the plasma membrane in p120ctn overexpressing cells was most likely due to the elevation of RhoA signalling. The reduction of Arf1 in the absence of p120ctn appears independent of RhoA and we suggest that it is caused by regulation of Arf1 recruitment and/or activation by p120ctn.

It was surprising that RhoGEF2-RNAi led to a uniform elevation of Arf1-GFP, as it affects RhoA signalling only at the AP borders (Bulgakova et al., 2013). Indeed, Arf1 was uniformly localized at the plasma membrane in all cases. FRAP of Arf1 at the plasma membrane demonstrated that it recovered almost completely within 25 sec (Fig. 5I), which indicated a highly dynamic exchange. Therefore, we suggest that the effect of RhoGEF2-RNAi on Arf1 at the DV borders is indirect: reduced RhoA signalling results in elevated recruitment of Arf1 at the AP borders, followed by rapid redistribution around the cell periphery and an overall elevation of Arf1-GFP signal at the cell surface. Overall, we identified two levels of p120ctn regulation of Arf1 activity and endocytosis: first, it either directly or indirectly promotes Arf1 membrane recruitment, and second, it indirectly inhibits this recruitment by promoting RhoA signalling activation.

### 6) Adhesion dynamics regulate cells shape

So far, we have demonstrated that p120ctn regulates cortical actin via RhoA signalling, and E-cad dynamics via both RhoA and Arf1. Next, we asked how this regulatory network contributes to the cell shape changes caused by altered p120ctn levels. Since changes in Arf1 activity did not affect RhoA signalling, we first examined how inhibition and hyperactivation of Arf1 alone affected cell shape. To explore this, we used Arf1 regulatory constructs to alter signalling output. First, we used a dominant negative (Arf1^DN^) expressed in the engrailed segment of the epidermis (Fig. 6A). Prolonged exposure to the dominant negative Arf1^DN^ construct resulted in small rounded cells (Fig. S4), with no surviving larvae, consistent with previous reports (Carvajal-Gonzalez et al., 2015), likely due to gross perturbation of post-Golgi protein transport causing cell death (Jian et al., 2010; Luchsinger et al., 2018). Therefore, we acutely induced expression of Arf1^DN^, using temperature sensitive GAL80^ts^. The cells which expressed the Arf1^DN^ had a reduced aspect ratio (p=0.003, Fig. 6A-B). This suggests that the reduction of endocytosis, and therefore stabilization of E-cad, is sufficient to reduce the cell’s aspect ratio. This is further supported by the reduced aspect ratio observed in other cases in which E-cad was stabilized: namely the expression of Rho^CA^ and the overexpression of p120ctn (p<0.0001, Fig. 6C-D, see Fig. 2 and 4).

**Figure 6.**
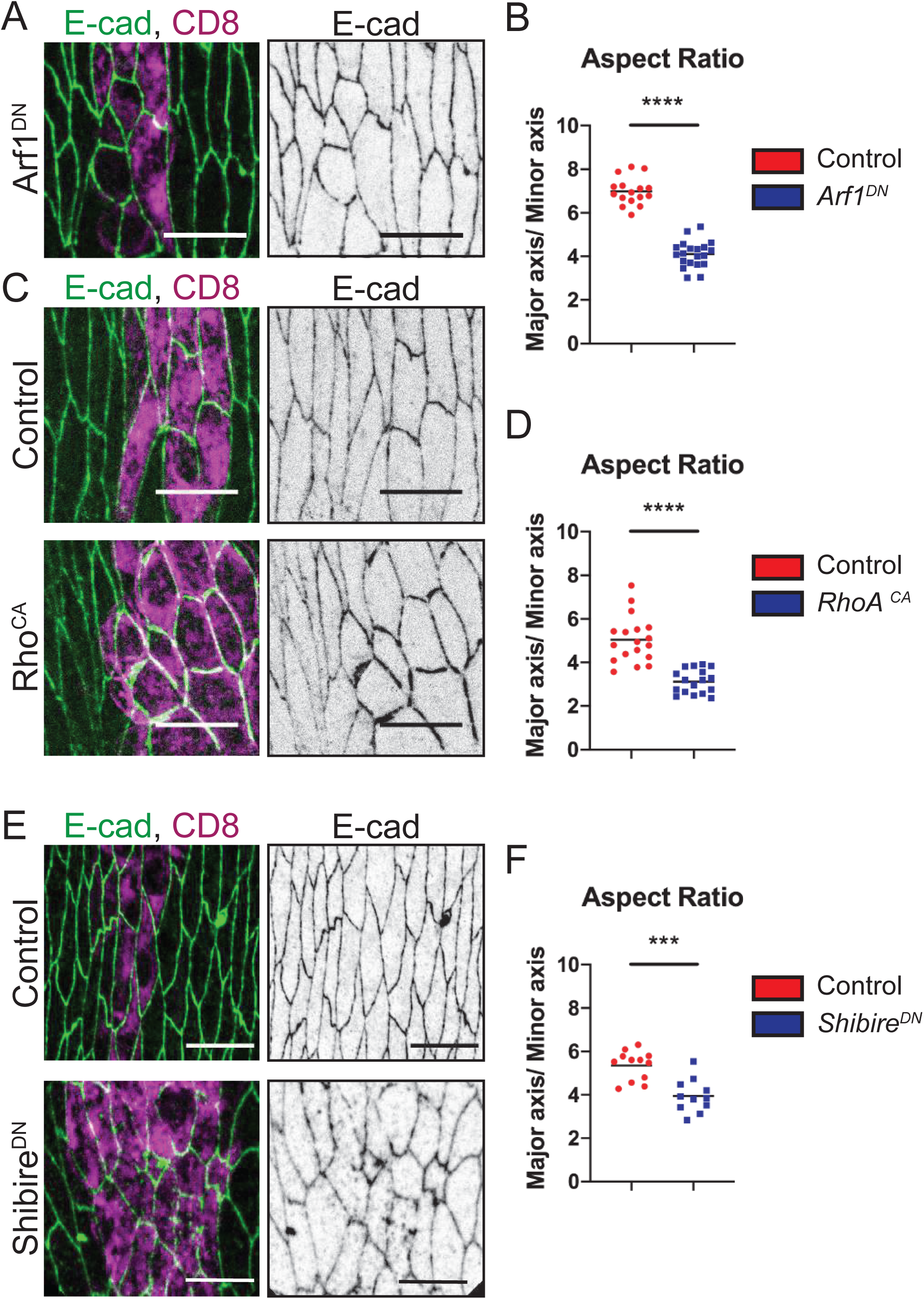
Inhibition of the dynamics of surface adhesion leads to reduced cell elongation. (**A-B**). Apical view (A) and cell aspect ratio (B) following downregulation of Arf1 (the induction of Arf1^DN^ expression for 4 hours). (**C-D**) Apical view (C) and cell aspect ratio (D) following upregulation of RhoA signalling (Rho^CA^). (**E-F**) Apical view (E) and cell aspect ratio (F) following inhibition of endocytosis by expression of the dominant-negative variant of dynamin (Shibire^DN^). Cell outlines are visualized by E-cad-GFP (A and C) or anti-E-cad antibody (E) in green (left) or grey (right). Cells expressing constructs are labelled with *UAS*::CD8-mCherry (A,C,E, magenta, left). ***, p < 0.001, and ****, p < 0.0001. All best-fit and membrane intensity data are in Table S1. 10-20 embryos per genotype were used for quantifications

To directly test if E-cad stabilization is sufficient to induce cell rounding, we used an alternative approach to inhibit E-cad endocytosis by overexpressing a dominant-negative dynamin Shibire (Shi^DN^) construct using *en*::GAL4. Expression of this construct stabilizes E-cad at the plasma membrane in a similar manner to the loss of p120ctn (Bulgakova et al., 2013). Indeed, cells expressing Shi^DN^ had reduced aspect ratio in comparison to control (p=0.0002, Fig. 6E-F). Overall, these results indicate that the dynamics of intercellular adhesion, mediated via endocytosis of E-cad, is a key factor determining the elongation of epidermal cells.

## Discussion

Epithelial cells *in vitro* are usually isotropic, and the application of external stretching or compressing forces induce an initial anisotropy in their shape (elongation), which is quickly resolved through cell rearrangements and divisions, or tissue three-dimensional deformation (Duda et al., 2019; Latorre et al., 2018; Nestor-Bergmann et al., 2019). By contrast, there are multiple examples of highly anisotropic elongated cells in whole organisms, including the epidermal cells used in this study but also in mammalian skin (Fig. 7A, Aw et al., 2016; Box et al., 2019). These elongated shapes are necessary for correct tissue and organism morphogenesis (Box et al., 2019; McCleery et al., 2019). In this study, we focused on the regulation of such elongated cell shape through the cross-talk between intercellular E-cad-mediated adhesion and cortical actin (the findings are summarized in Table 1). We provide *in vivo* evidence that p120ctn, a known regulator of E-cad dynamics and endocytosis (Bulgakova and Brown, 2016; Ireton et al., 2002; Nanes et al., 2012; Sato et al., 2011), mediates this cross-talk and regulates cell shape. It does so by promoting the activities of at least two small GTPases with opposing effects on E-cad dynamics: RhoA, which inhibits E-cad turnover, and Arf1, which promotes it (Fig. 7B). This is the first report of a role for Arf1 in the clathrin-mediated trafficking of E-cad from the membrane. We show a further interplay between these GTPases with RhoA preventing the localization of Arf1 at the plasma membrane (Fig. 7B). Finally, while p120ctn normally colocalizes with E-cad and is at higher levels on the DV borders, we show that it regulates RhoA activity only at the AP borders, which are under higher tension, suggesting a tension-dependent function for p120ctn.

**Figure 7.**
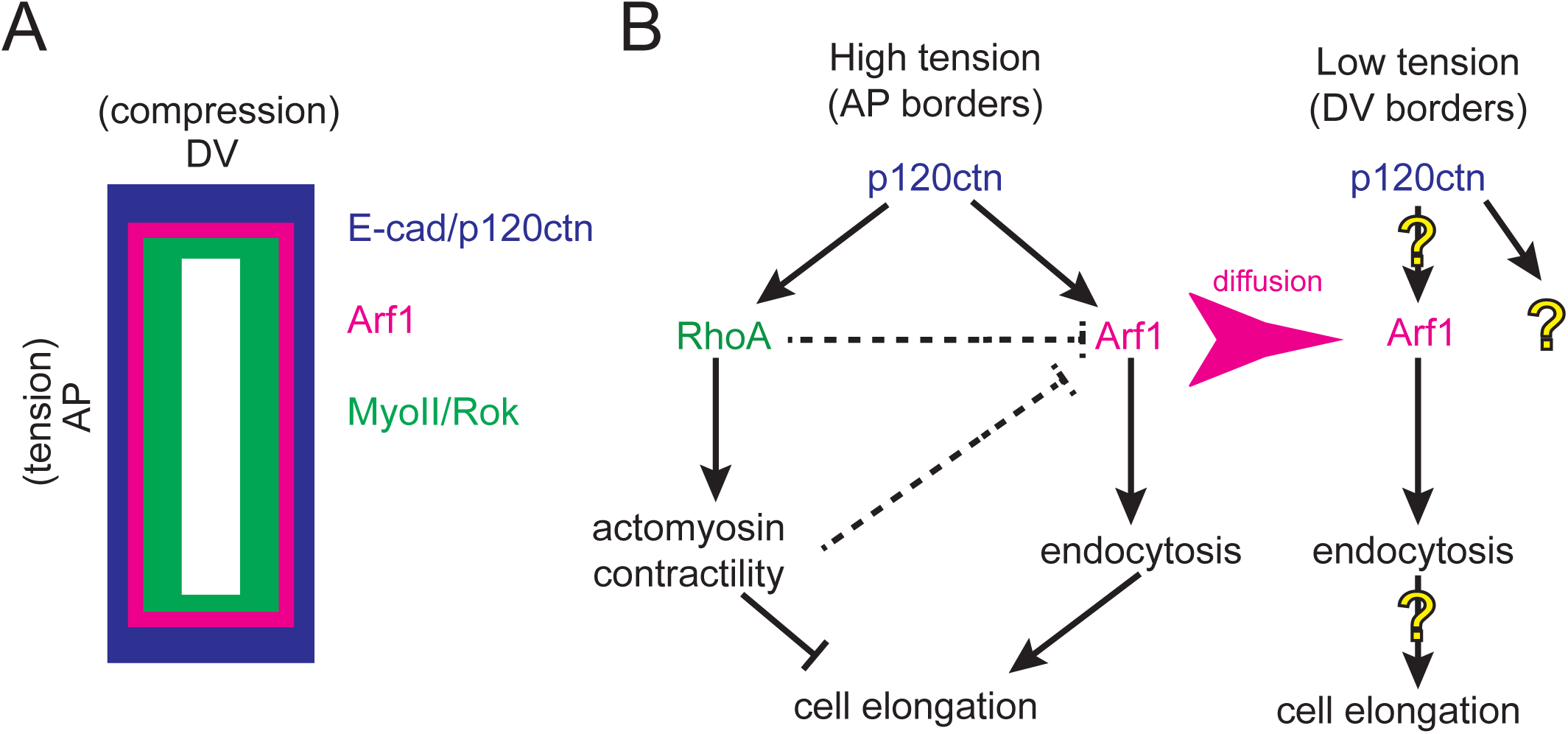
Model of cell shape regulation in anisotropic elongated cells in epidermis. **(A).** Schematic illustration of cellular anisotropy in embryonic epidermis of stage 15 *Drosophila* embryos. E-cad and p120ctn (blue) are enriched at the DV borders, whereas MyoII and Rok (green) at the AP borders, while Arf1 (magenta) is uniformly associated with the plasma membrane. The long AP borders are under tension, while the short DV borders are under compressive force (**B**). Model of cell shape regulation by cortical actin and E-cad dynamics at both cell borders. High cortical tension and actomyosin contractility at the long AP borders inhibit cell elongation, whereas adhesion dynamics, i.e. E-cad endocytosis, promote it. Actomyosin is activated downstream of RhoA signalling, while Arf1 GTPase promotes clathrin-mediated endocytosis. Furthermore, RhoA additionally inhibits this endocytosis by either directly or indirectly preventing Arf1 localization at the plasma membrane (dashed lines). Finally, both RhoA and Arf1 are activated downstream of p120ctn at the long AP borders. In contrast, at the short DV borders, where tension is low, RhoA seems to be low resulting in low MyoII and Rok levels and is thus not involved in cell elongation. It is also not regulated by p120ctn at these borders, despite p120ctn being normally enriched there. Arf1 promotes E-cad endocytosis at the DV borders. Arf1 is highly diffusive and its levels at the DV borders are most likely to be influenced by recruitment to the plasma membrane at the AP borders via diffusion (magenta arrow). Finally, it is unclear whether Arf1 or other molecules are regulated downstream of p120ctn at the DV borders, neither if E-cad dynamics via endocytosis contributes to cell shape and elongation (question marks).

**Table 1.** Summary of cell shape and protein distributions in the genotypes examined. Genotypes of the embryos presented in this study on the with corresponding changes observed in the aspect ratio of the cells; and the levels of representative cell adhesion, cytoskeletal, and signalling pathway components at both the AP and DV cell borders. OE – overexpression.

| Genotype | Aspect ratio | Amount at AP |  |  | Amount at DV |  |  |
| --- | --- | --- | --- | --- | --- | --- | --- |
|  |  | E-cad AP | MyoII AP | Arf1 AP | E-cad DV | MyoII DV | Arf1 DV |
| p120ctn OE | down | up | up | down | ns | ns | down |
| *p120ctn*<sup>-/-</sup> | ns | down | down | down | down | ns | down |
| *Rho*<sup>DN</sup> | ns | down | down | up | ns | down | up |
| *Rho*<sup>CA</sup> | down | ns | up | down | up | up | down |
| *Arf1*<sup>DN</sup> | down | ns | ns | down | ns | ns | down |
| *Arf1*<sup>CA</sup> | ns | ns | ns | up | ns | ns | up |
| RhoGEF2RNAi | ns | down | down | up | down | ns | up |
| *p120ctn*<sup>-/-</sup>; *Arf1*<sup>CA</sup> | down |  |  | up |  |  | up |
| *Shibire*<sup>DN</sup> | down | up |  |  | up |  |  |

Multiple studies have revealed that intercellular adhesions are in a state of constant remodelling through the processes of endocytosis and recycling (recently reviewed in Bruser and Bogdan, 2017). This recycling is important for junctional remodelling and cell rearrangements during morphogenesis (Le Droguen et al., 2015; Levayer et al., 2011a; Ogata et al., 2007; Sumi et al., 2018). At the same time, the contribution of adhesion dynamics, in particular E-cad recycling, to the shape of cells is largely overlooked, while the roles of cortical actin and the levels of intercellular adhesion in shaping cells are well established (Lecuit and Lenne, 2007). Here, we show that the stabilization of E-cad and thus inhibition of junctional remodelling prevents the elongation of the cells in *Drosophila* embryonic epidermis (Fig. 7B). This is supported by several lines of evidence: expression of Arf1^DN^, Shi^DN^, and Rho^CA^, all of which inhibit E-cad dynamic behaviour, causing loss of elongation. Moreover, the changes in cell elongation were observed only in the cases when E-cad was stabilized, suggesting a central role for E-cad dynamics in cell elongation. However, it is not the only contributing factor: in *p120ctn* mutant embryos cell elongation was not affected despite E-cad being stabilized. As E-cad stabilization in these embryos was accompanied by a reduction of RhoA signalling at the AP borders and a uniform reduction in E-cad junctional levels, either one or both of these factors are likely to counteract the stabilization of E-cad resulting in normal elongation. We speculate that RhoA activity and cortical actin organization are likely to be responsible for compensating for E-cad stabilization (Fig. 7B). The contractility of cortical actin normally acts to reduce the surface length whereas the levels of adhesion acts oppositely (Lecuit and Lenne, 2007), thus reduced MyoII in the *p120ctn* mutants would act to increase the length of the AP borders, whereas reduced E-cad would act to reduce it. Indeed, the strongest reduction in cell elongation was observed upon the expression of Rho^CA^, which stabilized E-cad without changing its levels at the AP borders and promoted actomyosin contractility. Therefore, we propose that cell elongation is the product of the counteraction of E-cad dynamics and cortical contractility.

However, due to the tight cross-regulation of E-cad adhesion, cortical actin, and endocytic machinery it is impossible to affect one without affecting the others. This makes it difficult to dissect the exact contribution of each component. For example, although the expression of Arf1^DN^ did not have any obvious effect on Rok localization, we cannot exclude the possibility that it altered the organization of cortical actin in another way. Indeed, Arf1 regulates actin assembly at the Golgi by acting both upstream and downstream of Cdc42 (Myers and Casanova, 2008). Similarly, dynamin, which is a large GTPase, regulates actin in GTPase-dependent manner in addition to driving endocytosis (Gu et al., 2010; Lee and De Camilli, 2002). Furthermore, the inhibition of E-cad endocytosis using a dominant-negative variant of the GTPase Rab5 resulted in an elevated level of actin in epidermal cells in *Drosophila* embryos (Mateus et al., 2011). Despite this difficulty in distinguishing the direct effects of endocytosis from indirect effects through the regulation of the cortex, our results establish that endocytosis is an important factor in regulating cell shape.

While the roles of RhoA and p120ctn in E-cad endocytosis are long-established (Davis et al., 2003; Ellis and Mellor, 2000), the Arf1 dependent recruitment of clathrin had only been shown to occur at the Golgi by recruiting the Adaptor 1 protein (Carvajal-Gonzalez et al., 2015). The function of Arf1 at the plasma membrane has been described in dynamin-independent endocytosis, which was presumed to be clathrin-independent (Kumari and Mayor, 2008). However, the involvement of Arf1 in multiple endocytic pathways has not been fully explored. We have shown that Arf1’s capacity to recruit clathrin is exploited by p120ctn to facilitate the endocytosis of E-cad. Whether this requires AP2, a well-documented plasma membrane resident clathrin adaptor, has yet to be determined. Our findings provide a mechanistic insight into the pro-endocytic activity of p120ctn which has only recently come to light (Bulgakova and Brown, 2016; Sato et al., 2011) and elaborates the number of known p120ctn interactors. The activities of Arf1 and RhoA are antagonistic, which was also seen in previous studies on the cellularization of the early embryo (Lee and Harris, 2013).

In contrast to Arf1, the regulation of RhoA by p120ctn has been shown in many studies, although the exact effects and mechanisms seem to be context-dependent (Anastasiadis et al., 2000; Derksen and van de Ven, 2017; Lang et al., 2014; Taulet et al., 2009; Yu et al., 2016; Zebda et al., 2013). We demonstrate that in the epidermal cells in *Drosophila* embryos, p120ctn leads to activation of RhoA specifically at the AP but not DV borders (Fig. 7B). At the same time, we show that p120ctn loss uniformly reduces recruitment of Arf1 to the plasma membrane. However, we also find that Arf1 is very dynamic and the changes at the DV borders, for example upon the downregulation of RhoGEF2, are likely to be an indirect consequence of the effect at the AP borders (Fig. 7B). It is therefore currently unclear if p120ctn performs any direct activity at the DV borders, or promotes RhoA and Arf1 signalling only at the high-tension AP borders. Indeed, recent evidence has suggested p120ctn has mechanosensing properties (Iyer et al., 2019). As we show that the AP borders are at higher tension than the DV, our data further suggest that p120ctn has the properties of a mechanotransducer. Additionally, using laser ablation we demonstrated that p120ctn directly modulates tension at these AP borders, providing further evidence for the novel role of p120ctn in mechanotransduction. It is yet to be determined if p120ctn is directly sensing the tension though a conformational change similarly to other components of cell-cell adhesion such as α-catenin and vinculin (Bays and DeMali, 2017; Yao et al., 2014), or is regulated by another mechanotransducer. Our findings expand the ever-increasing list of proteins involved in mechanotransduction and mechanical cell responses.

Altogether, our findings demonstrate that cell elongation in a tissue is regulated through the opposing action of cortical actin contractility and adhesion endocytosis, which are however closely interconnected and regulate each other. Adhesion components modify cortical actin while the latter alters adhesion endocytosis. Considering that all of the molecules studied are expressed in all epithelia across evolution, we speculate that this system is likely to be more broadly applicable in development and a general feature of cell biology.

## Materials and Methods

### Fly stocks and genetics

Flies were raised on standard medium. The GAL4/UAS system (Brand and Perrimon, 1993) was used for all the specific spatial and temporal expression of transgenic and RNAi experiments. The GAL4 expressional driver used for all experiments was *engrailed::GAL4* (*en*::GAL4, Bloomington number 30564). The following fly stocks were used in this study (Bloomington number included where applicable): E-cad-GFP (*shg::E-cadherin-GFP*) (60584), Shg-Cherry (*shg::mCherry*, 59014), *UAS*::CLC-GFP (7109), *UAS*::Arf1-GFP (gift from T. Harris), Zipper-YFP (*Myosin II-*YFP, Kyoto Stock Center 115082), *sqh*::Rok^K116A^-Venus (gift from J. Zallen), *UAS*::Arf1-T31N (DN) and *UAS::*Arf1-Q71L (CA) (Dottermusch-Heidel et al., 2012), *UAS*::Rho1-N19 (DN) (7328), *tubulin*::GAL80^TS^ (7017), *ubi*::p120ctn-GFP (7190), *UAS*::p120ctn-GFP (7192), and *UAS*::Shibire^K44A^ (*UAS*::Shi-DN, 5822). The p120ctn mutant genotype (p120ctn^308^/ Δp120) used was derived from crossing two deficiency stocks: homozygously viable p120ctn^308^ females (Myster et al., 2003) with homozygously lethal Df(2R)M41A8/ CyO, *twi::*Gal4, *UAS::*GFP males (740). Thus, the p120ctn mutants examined lacked both maternal and zygotic contributions.

### Embryo collection and fixation

Embryos were collected at 25°C at 3-hour time intervals and allowed to develop at 18°C for 21 hours to reach the desired developmental stage, except for the acute induction experiments, where embryos were allowed to develop for 13 hours at 18°C and shifted to 29°C for 4 hours. Then embryos were dechorionated using 50% sodium hypochlorite (bleach, Invitrogen) in water for 4 minutes, and extensively washed with deionized water prior to fixation. Fixation was performed with a 1:1 solution of 4% formaldehyde (Sigma) in PBS (Phosphate Buffered Saline) and heptane (Sigma) for 20 minutes on an orbital shaker at room temperature. Embryos were then devitellinized in 1:1 solution of methanol and heptane for 20 sec with vigorous agitation. Following subsequent methanol washes the fixed embryo specimens were stored at −20°C in methanol until required.

### Embryo live imaging

Embryos were collected and dechorionated as described above. Once washed with deionized water embryos were transferred to apple juice agar segments upon microscope slide. Correct genotypes were selected under a fluorescent microscope (Leica) using a needle. Embryos were positioned and orientated in a row consisting of 6-10 embryos per genotype. Following this, embryos were transferred to pre-prepared microscope slides with Scotch tape and embedded in Halocarbon oil 27 (Sigma). Embryos were left to aerate for 10 minutes prior to covering with a cover slip and imaging.

For laser ablation, following orientation and positioning the embryos were transferred to a 60mm x 22mm coverslip which had been pre-prepared by applying 10 µl of Heptane glue along a strip in the middle of the coverslip orientated with the long axis. The coverslip was attached to a metal slide cassette (Zeiss), and the embryos were embedded in Halocarbon oil 27 before imaging.

### Molecular cloning

The p120ctn full length cDNA was obtained from Berkeley Drosophila Genome Project (BDGP), supplied in a pBSSK vector. This was sub-cloned into a (pUAS-k10.attB) plasmid using standard restriction digestion with NotI and BamHI (New England Biolabs) followed by ligation with T4 DNA ligase (New England Biolabs) and transformation into DH5a competent *E.coli* cells (Thermo Fisher Scientific). Prior to injection plasmids were test digested and sequenced (Core Genomic Facility, University of Sheffield). Plasmids were prepared for injection using standard miniprep extraction (Qiagen) and submitted for injection (Microinjection service, Department of Genetics, University of Cambridge) into the attP-86Fb stock (Bloomington stock 24749). Successful incorporation of the transgene was determined by screening for (*w^+^*) in the F1 progeny.

### Immunostaining

The embryos were washed three times in 1 ml of PBST (PBS with 0.05% Triton) with gentle rocking. Blocking of the embryos prior to staining was done in 300 µl of a 1% NGS (Normal Goat Serum) in PBST for 1 hour at room temperature with gentle rocking. For staining the blocking solution was removed, 300 µl of the primary antibody: 1:100 rat anti-E-cad (DCAD2, DSHB), 1:10 mouse anti-engrailed (4D9, DSHB), or 1:500 anti-Golgi (Golgin-245, Calbiochem) in fresh blocking solution was added and the embryos were incubated overnight at 4°C with orbital rotation. Then, embryos were washed three times with 1 ml of PBST. A 300 µl 1:300 dilution of the secondary antibody (goat Cy3-or Cy5-conjugated IgG, Invitrogen) was added, and the embryos incubated either overnight at 4°C with orbital rotation or for 2 hours at room temperature. Then embryos were washed three time with PBST, following which they were incubated with 50-70 µl of Vectashield (Vector Laboratories) and allowed to equilibrate for a period of 2 hours before being mounted on microscope slides (Thermo).

### Microscopy, data acquisition and FRAP

All experiments except for laser ablation were performed using an up-right Olympus FV1000 confocal microscope with a 60x/1.40 NA oil immersion objective. Shi^DN^ expressing embryos and the corresponding control were imaged using 100x/1.40 NA UPlanSApo objective. All measurements were made on dorsolateral epidermal cells of embryos, which were near or just after completion of dorsal closure, corresponding to the end of Stage 15 of embryogenesis. For fixed samples 16-bit images were taken at a magnification of 0.051µm/pixel (1024×1024 pixel XY-image) or 0.062 µm/pixel (Shi^DN^ embryos and the corresponding control) with a pixel dwell of 4µm/pixel. For each embryo, a Z-axis sectional stack through the plane of the AJs was taken, which consisted of six sections with a 0.38 µm intersectional spacing. The images were saved in the Olympus binary image format for further processing.

For E-cad FRAP (adapted from Bulgakova et al., 2013) 16-bit images were taken at a magnification of 0.093 µm/pixel (320×320 pixel XY-image). In each embryo, several circular regions of 1 µm radius were photobleached at either DV or AP junctions resulting in one bleach event per cell. Photobleaching was performed with 8 scans at 2 µs/pixel at 50-70% 488 nm laser power, resulting in the reduction of E-cad-GFP signal by 60–80%. A stack of 6 z-sections spaced by 0.38 µm was imaged just before photobleaching, and immediately after photobleaching, and then at 20 s intervals, for a total of 15 minutes.

As rate of endocytosis depends on external factors, such as temperature [103], controls were examined in parallel with experimental conditions in all experiments with CLC-GFP. This dependence might also account for slight variation between datasets, e.g. compare Fig. 2A and 2G. For CLC-GFP FRAP, 16-bit images were taken at a magnification of 0.051µm/pixel (256×256 pixel XY-image). In each embryo a single plane was selected in centre of the AJ band using Shg-Cherry for positioning. An area encompassing a transverse region orthogonal to the axis of the engrailed expressing cells was selected (140×60 pixels) was photobleached with 1 scan at 2 µm/pixel using 100% 488nm laser power resulting in reduction of CLC-GFP signal by 70-80%. Images were taken using continuous acquisition at a frame rate of 2 sec^-1^. Prior to bleaching a sequence of 10 images was taken, and a total of 400 frames corresponding to 3.5 minutes were taken.

### Data processing and statistical analysis

*Membrane intensity and cell shape*: Images were processed in Fiji (https://fiji.sc) by generating average intensity projections of the channel required for quantification. Masks were created by processing background-subtracted maximum intensity projections using the Tissue Analyzer plugin in Fiji (Aigouy et al., 2016). Quantification of the membrane intensity at the AP and DV borders and cell elongation (aspect ratio) was done as described previously using a custom-built Matlab script (Bulgakova and Brown, 2016) found at (https://github.com/nbul/Intensity). In short, cells were identified as individual objects using the created masks, and their eccentricities were calculated. The aspect ratio was calculated from the eccentricity as 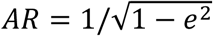, where *e* is eccentricity. At the same time, the individual borders were identified as objects by subtracting a dilated mask of vertices from a dilated mask of cell outlines. The mean intensity and orientation of each border were calculated. The average border intensities (0-10° for the AP and 40-90° for the DV borders relatively to cell mean orientation) were calculated for each embryo and used as individual data points to compare datasets. The average cytoplasmic intensity was used for the background subtraction. In the case of quantification of Arf1-GFP membrane intensity, due to high Arf1-GFP presence inside cells both in Golgi and cytoplasm, the mean intensity of embryonic areas not expressing the GFP-tagged transgene were used for background subtraction. Statistical analysis was performed in Graphpad Prism (https://www.graphpad.com/scientific-software/prism/). First, the data was cleaned using ROUT detection of outliers in Prism followed by testing for normal distribution (D’Agostino & Pearson normality test). Then, the significance for parametric data was tested by either a two-way ANOVA or two-tailed t-test with Welch’s correction. *E-cad FRAP*: images were processed by using the grouped Z-projector plugin in Fiji to generate average intensity projections for each time-point. Following this the bleached ROI, control ROI and background intensity were manual measured for each time point. This data was processed in Microsoft Excel. First the intensity of the bleached ROI at each time point was background subtracted and normalized as following: *I* _*n*_ = (*F*_*n*_ − *BG*_*n*_)⁄(*FC*_*n*_ − *BG*_*n*_), where *F_n_* – intensity of the bleached ROI at the time point *n*, *FC_n_* – intensity of the control unbleached ROI of the same size at the plasma membrane at the time point *n*, and *BG_n_* – background intensity, measured with the same size ROI in cytoplasm at the time point *n*. Than the relative recovery at each time point was calculated using the following formula: *R*_*n*_ = (*I*_*n*_ − *I*_1_)⁄(*I*_0_ − *I*_1_), where *I_n_*, *I*_1_ and *I*_0_ are normalized intensities of bleached ROI and time point *n*, immediately after photobleaching, and before photobleaching respectively. These values were input to Prism and nonlinear regression analysis was performed to test for best fit model and if recoveries were significantly different between cell borders or genotypes. The recovery was fit to either single exponential model in a form of *f*(*t*) = 1 − *F*_*im*_ − *A*_1_*e*^−*t*⁄*T_fast_*^, or to bi-exponential model in a form of *f*(*t*) = 1 − *F*_*im*_ − *A*_1_*e*^−*t*⁄*T_fast_*^ − *A*_2_*e*^−*t*⁄*T_slow_*^, where *F_im_* is a size of the immobile fraction, *T_fast_*and *T_slow_* are the half times, and *A*_1_ and *A*_2_ are amplitudes of the fast and slow components of the recovery. An F-test was used to choose the model and compare datasets.

*CLC-GFP FRAP:* measurements of all intensities, i.e. the bleached ROI, control ROI and the background, and normalization were done using a custom-build Matlab script (http://github.com/nbul/FRAP) using the same algorithm as described for E-cad FRAP. Curve fitting and statistical analysis was performed in Graphpad Prism using a nonlinear regression analysis as described for E-cad FRAP.

*Sample size:* Each experiment was performed on multiple embryos, which represent biological replicates. For each embryo, 2-3 ROIs were selected for E-cad FRAP analysis, and these were treated as technical replicates and were averaged per wing. The minimum sample size was estimated based on the mean values and the standard deviation of a control set of embryos to allow detection of differences of 20% in the means, in a pair-wise comparison, with a power of 0.8 and α 0.05 (using http://powerandsamplesize.com/, 9 embryos for fluorescence intensity comparison and aspect ratio, and 5 embryos for FRAP). As standard deviations were larger for some genotypes, we aimed for 15 and 6 embryos per genotype, respectively. The exact numbers of embryos used in each experiment are in Table S1.

### Laser Ablation

Nanoablation of single junctions was performed to provide a measure of junctional tension. Embryos were imaged on a Zeiss LSM 880 microscope with an Airyscan detector, an 8-bit image at 0.053 µm/pixel (512×512 pixel XY-Image) resolution with a 63x objective (1.4 NA) at 5x zoom and 2x averaging was used. An illumination wavelength of 488 nm and 0.5% laser power were used. Images were captured with a 0.5 µm z-spacing. Narrow rectangular ROIs were drawn across the centre of single junctions and this region was ablated using a pulsed TiSa laser (Chameleon), tuned to 760 nm at 45% power. Embryos were imaged continuously in a z-stack consisting of 3 z-slices. The initial recoil rate of vertices at the ends of ablated junctions was quantified by measuring the change in distance between the vertices and dividing by the initial time step. Statistical analysis was performed in Graphpad Prism: using a two-tailed t-test with Welch’s correction.

## Acknowledgments

The authors first wish to thank Rob Tetley and Yanlan Mao for assistance with laser ablation experiments. The authors also with to acknowledge the advice and guidance of Professor David Strutt, University of Sheffield, and his lab. The authors thank to the technical staff of the Wolfson Light Microscopy Facility (LMF) and the Fly Facility, the University of Sheffield, without whom this work would not be possible. This work was supported by grant BB/P007503/1 from the UK Biotechnology and Biological Sciences Research Council.

## Author Contribution

J.G and N.A.B. designed and performed experiments and wrote the manuscript.

## Supplementary Information

**Table S1. Numerical values for each experiment presented in paper.**

**Figure S1.**
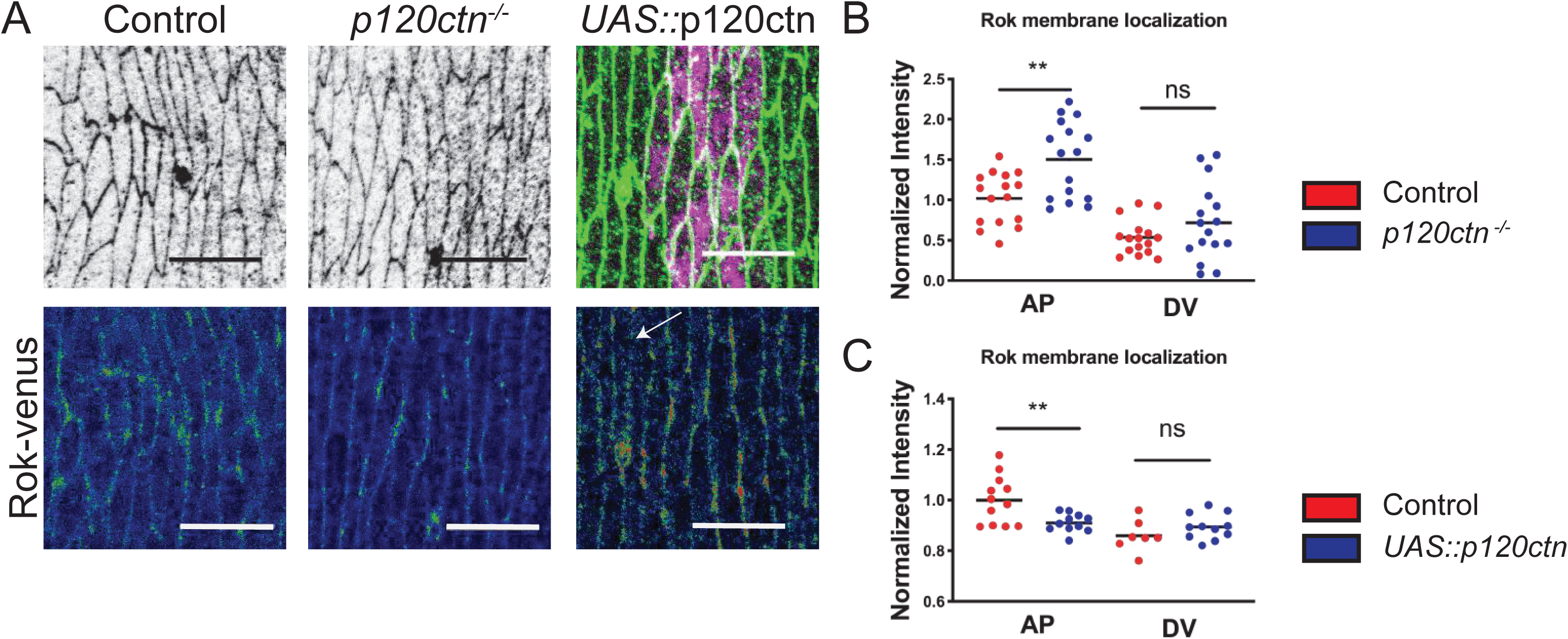
Rok-Venus is effected is the same manner as MyoII by changes the levels of p120ctn. **A.** Apical view of the epidermis of control, *p120ctn^-/-^* mutant, and p120ctn overexpression (*UAS::*p120ctn) embryos visualized with anti-E-cad antibody (**A**, grey in the top images, green in the top right image), *UAS*::CD8-mCherry (**A**, magenta in the right image, marks cells expressing *UAS::*p120ctn) and Rho-Kinase (Rok) tagged with Venus (**A,** lower panel rainbow pixel intensity spectrum, membrane indicated by arrow). **B-C.** The amount of Rok-Venus in the *p120ctn^-/-^* mutant (**B**) and p120ctn overexpressing embryos (**C**). Scale bars – 10 µm. Statistical analysis between cell borders measured by two-way ANOVA. **, p < 0.01. 15-20 embryos per genotype were used for quantifications.

**Figure S2.**
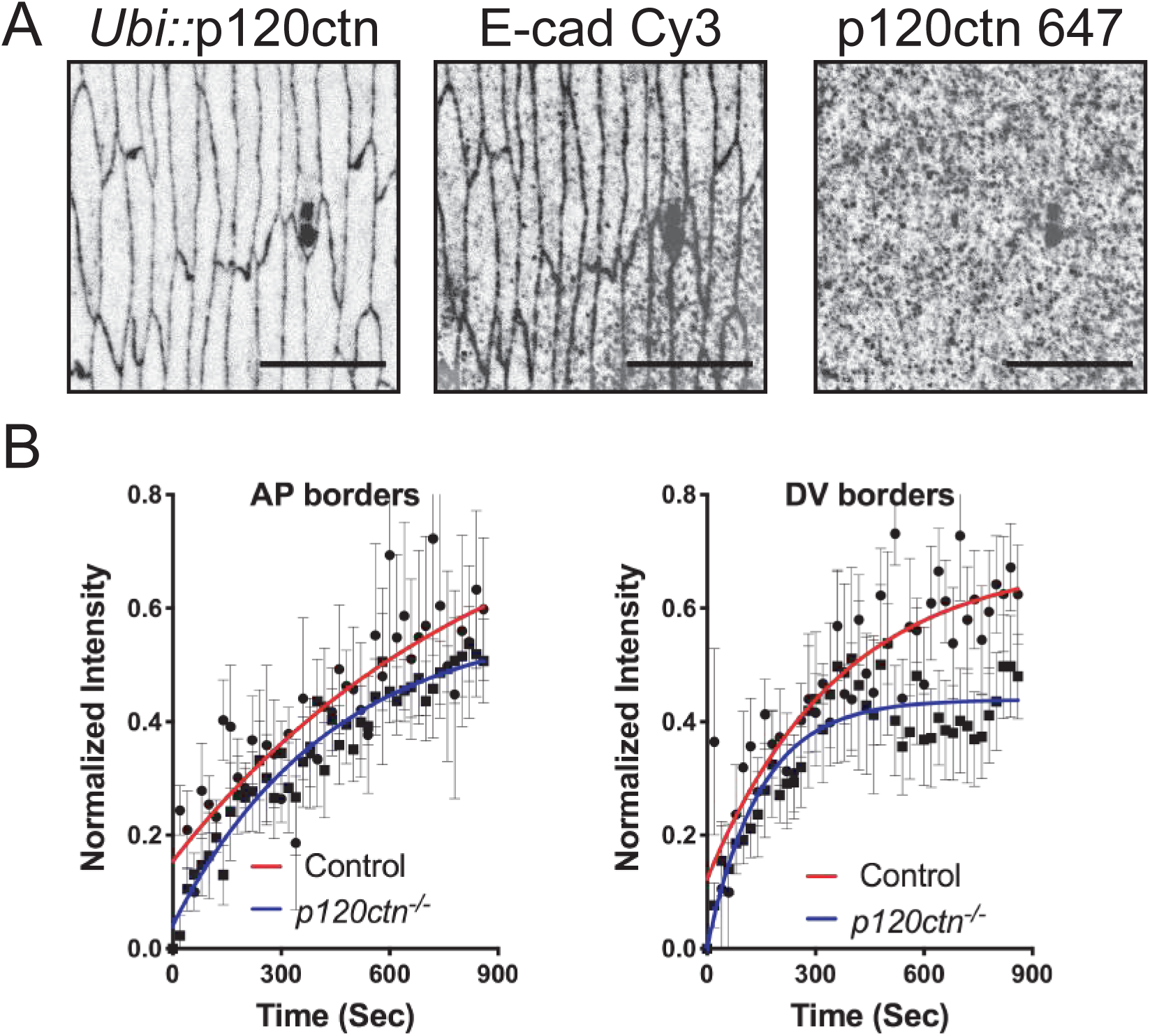
p120ctn antibody staining fails to replicate localization of GFP tagged variant and genetic ablation results in stabilization of endogenously tagged E-cad-GFP. **A.** Apical view of epidermal cells of the *ubi::*p120ctn-GFP expressing embryos. The localization of the p120ctn-GFP (grey, left panel) compared to the localisation of E-cad stained with an antibody (grey, middle panel) the staining pattern of the anti-p120ctn antibody in the same cells (grey, right panel). **B.** The FRAP curves of endogenously tagged E-cad-GFP in *p120ctn^-/-^* mutant embryos. AP borders (left graph) and DV borders (right graph) are shown separately. All best-fit and membrane intensity data are in Table S1. Scale bars – 10 µm. 8-10 embryos used for quantification in FRAP.

**Figure S3.**
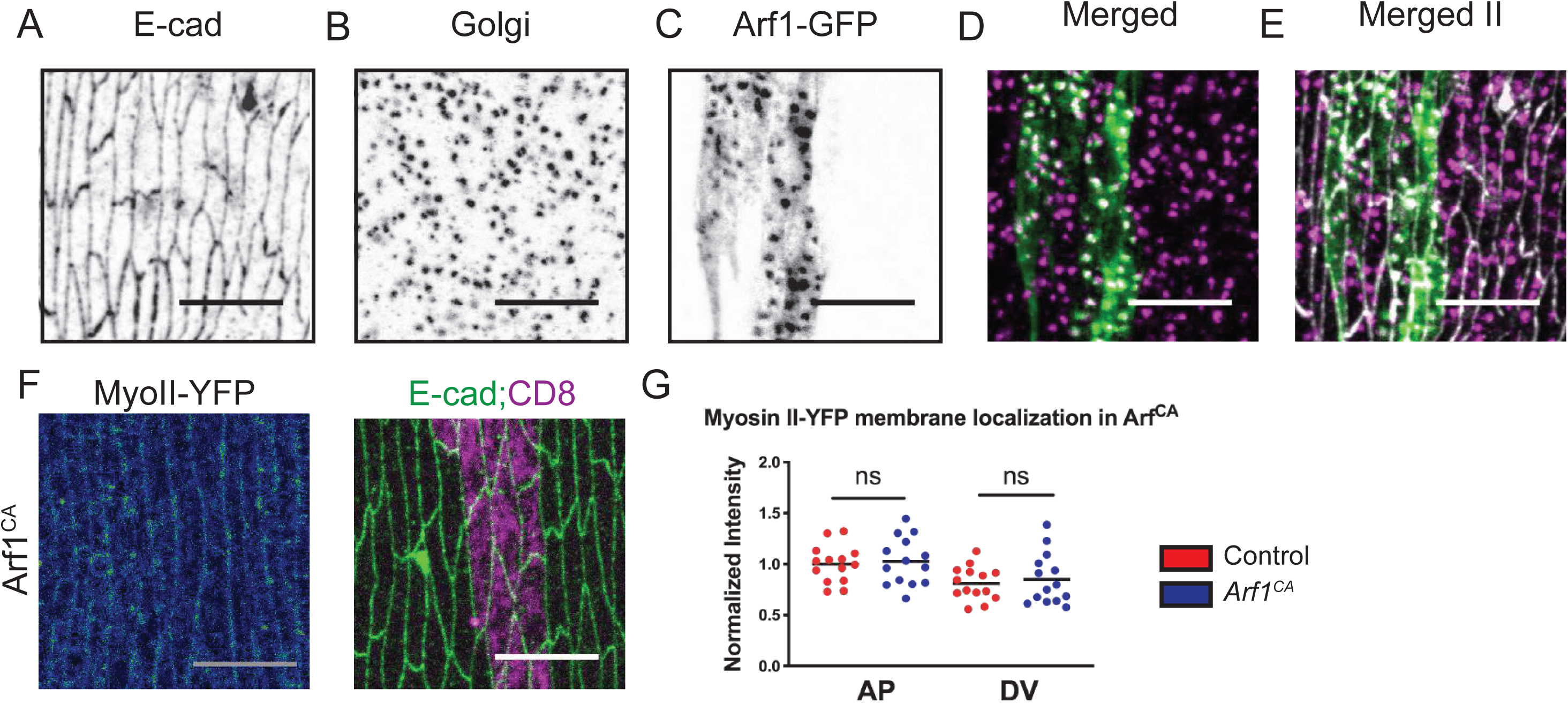
Arf1-GFP cytoplasmic puncta are representative of the Golgi apparatus and hyperactivation of Arf1 signalling has no effect on MyoII-YFP. **A-E.** Apical view of the dorsolateral epidermis of *UAS::*Arf1-GFP expressing control embryo with cells borders visualized with anti-E-cad antibody (black in **A**, white in **E**), Golgi marked by immunostaining of Trans-Golgi (black in **B**, magenta in D and E), and Arf1-GFP in the *engrailed* expressing cells (black in **C,** green in **D** and **E)**. The same region is shown in main text (see Fig. 5). (**F-G**) Localization of MyoII-YFP (rainbow in left image) in the cells expressing a constitutively active Arf1 (Arf1^CA^) with *en*::GAL4 marked by *UAS::*CD8-Cherry (magenta in right image). Cell borders are visualized with anti-E-cad antibody staining (green in right image). **G.** The amount of MyoII-YFP at the plasma membranes of the Arf1^CA^ cells and the adjacent internal control cells. Scale bar is 10 µm. Statistical analysis between cell borders measured by two-way ANOVA. 15-20 embryos were used for quantification.

**Figure S4.**
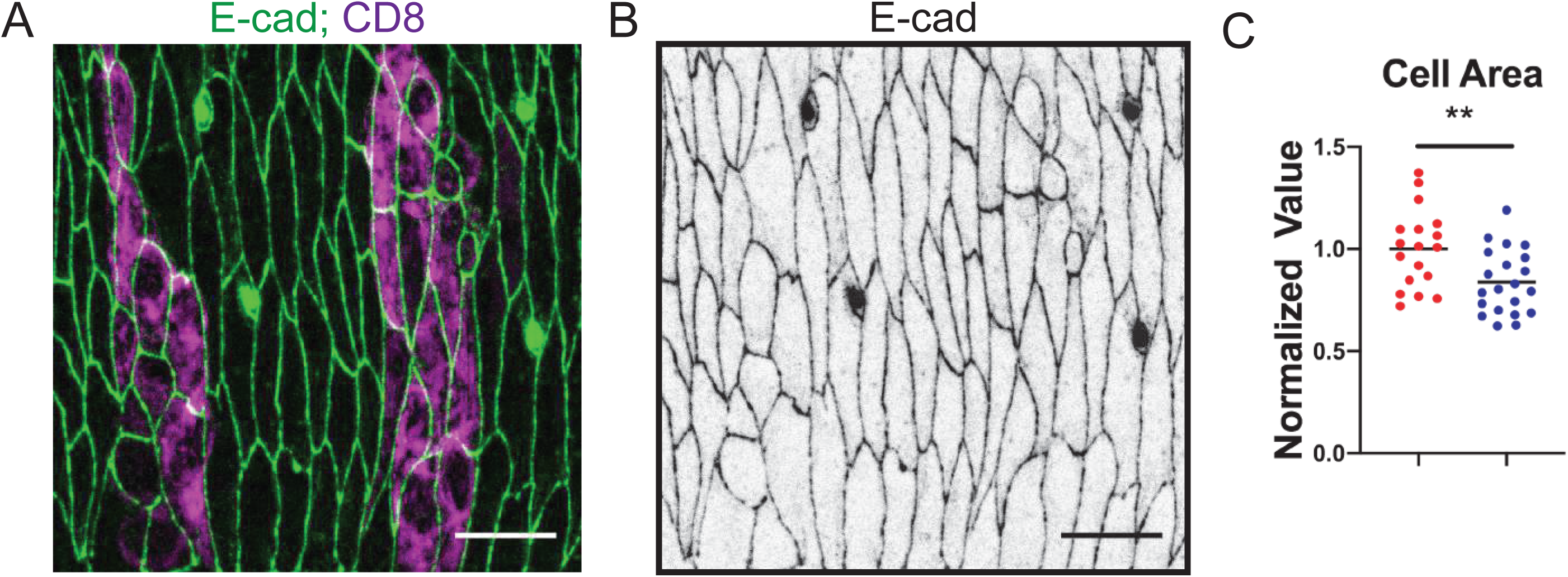
Arf1^DN^ induces cell morphology changes leading to cell death. **A-B.** Apical view of the dorsolateral epidermis of Arf1^DN^ transgene expressing embryos, co-expressing *UAS::*CD8-cherry in the engrailed stripes to mark the cells (magenta in **A**). The cell borders are marked by Shg-GFP (green in A, black in B). Scale bar is 10 µm. **C.** Cell area of the internal control and Arf1^DN^ expressing cells, normalized to control. **, p < 0.01.

